# Single-cell analysis uncovers differential regulation of lung γδ T cell subsets by the co-inhibitory molecules, PD-1 and TIM-3

**DOI:** 10.1101/2021.07.04.451035

**Authors:** Sarah C. Edwards, Ann Hedley, Wilma H. M. Hoevenaar, Teresa Glauner, Robert Wiesheu, Anna Kilbey, Robin Shaw, Katerina Boufea, Nizar Batada, Karen Blyth, Crispin Miller, Kristina Kirschner, Seth B. Coffelt

**Affiliations:** Cancer Research UK Beatson Institute, Glasgow, UK; Institute of Cancer Sciences, University of Glasgow, UK; Institute of Genetics and Molecular Medicine, University of Edinburgh, Edinburgh, UK

**Author notes:** Corresponding author Seth B. Coffelt Cancer Research UK Beatson Institute, Garscube Estate, Switchback Road, Glasgow, United Kingdom, G61 1BD.

## Abstract

IL-17A-producing γδ T cells within the lung consist of both Vγ6^+^ tissue-resident cells and Vγ4^+^ circulating cells that play important roles in homeostasis, inflammation, infection, tumor progression and metastasis. How these γδ T cell subsets are regulated in the lung environment during homeostasis and cancer remains poorly understood. Using single-cell RNA sequencing and flow cytometry, we show that lung Vγ6^+^ cells express a repertoire of cell surface molecules distinctive from Vγ4^+^ cells, including PD-1 and ICOS. We found that PD-1 functions as a co-inhibitory molecule on Vγ6^+^ cells to reduce IL-17A production, whereas manipulation of ICOS signaling fails to affect IL-17A in Vγ6^+^ cells. In a mammary tumor model, ICOS and PD-1 expression on lung Vγ6^+^ cells remained stable. However, Vγ6^+^ and Vγ4^+^ cells within the lung pre-metastatic niche increased expression of IL-17A, IL-17F, amphiregulin (AREG) and TIM-3 in response to tumor-derived IL-1β and IL-23, where the upregulation of TIM-3 was specific to Vγ4^+^ cells. Inhibition of either PD-1 or TIM-3 in mammary tumor-bearing mice further increased IL-17A by Vγ6^+^ and Vγ4^+^ cells, indicating that both PD-1 and TIM-3 function as negative regulators of IL-17A-producing γδ T cell subsets. Together, these data demonstrate how lung γδ T cell subsets are differentially controlled by co-inhibitory molecules in steady-state and cancer.

## INTRODUCTION

Mouse IL-17A-producing γδ T cells predominantly express Vγ4 or Vγ6 T cell receptor (TCR) chains (O’Brien and Born, 2020). In adult mice, Vγ4^+^ and Vγ6^+^ cells are self-renewing and persist for a long time in tissues (Sandrock et al., 2018). Functionally, these two subsets are indistinguishable. Their phenotype is almost identical, but there are some differences in the molecules they express, their behavior and their distribution (O’Brien and Born, 2020). For example, the TCR repertoire for the Vγ6 population utilises an invariant clone with few semi-invariant clones for the Vγ6Vδ1 pairing (Sandrock et al., 2018; Wei et al., 2015). The IL-17A- producing Vγ4^+^ cells comprise several semi-invariant oligoclonal TCRs that pair with δ4 or δ5 chains (Chen et al., 2019; Kashani et al., 2015; Wei et al., 2015). TCR expression levels are different between the two subsets with Vγ6^+^ cells expressing higher levels of CD3 subunits with their TCR (Paget et al., 2015). Vγ6^+^ cells are enriched in mucosal tissues such as the lung, dermis, and uterus as tissue resident cells (Monin et al., 2020; Sandrock et al., 2018; Tan et al., 2019); they are scarcely found in lymphoid organs of young mice, but accumulate in lymph nodes as mice age (Chen et al., 2019). Vγ4^+^ cells are more migratory than Vγ6^+^ cells, endowed with the ability to traffic from tissue to lymph nodes (McKenzie et al., 2017; Ramirez-Valle et al., 2015). In addition to IL-17A, these subsets can produce several other cytokines, such as IL-17F, IL-22, as well as the epidermal growth factor (EGF) receptor ligand, amphiregulin (AREG) (Jin et al., 2019; Sutton et al., 2009; Tan et al., 2019). Both Vγ4^+^ and Vγ6^+^ cells express the cell surface molecules, CCR6, CCR2, CD44, IL-23R, IL-1R1, and IL-7Rα (Haas et al., 2009; McKenzie et al., 2017; Michel et al., 2012; Ribot et al., 2009; Sutton et al., 2009; Tan et al., 2019); the transcription factors, RORγt and MAF (Zuberbuehler et al., 2019); as well as B lymphoid kinase (BLK) (Laird et al., 2010). Vγ4^+^ cells express SCART2 and CD9, whereas Vγ6^+^ cells express SCART1 and PD-1 (Kisielow et al., 2008; Tan et al., 2019). During development, SOX4 and SOX13 transcription factors regulate the genesis of Vγ4^+^ cells (Gray et al., 2013; Malhotra et al., 2013), while PLZF is required for Vγ6^+^ cells (Kreslavsky et al., 2009; Lu et al., 2015). Vγ6^+^ cells further differ from V 4β^+^ cells in their unique regulation by β2 integrins and BCL-2 family members (McIntyre et al., 2020; Tan et al., 2019). Although several studies have shed light on the similarities and differences between Vγ4^+^ and Vγ6^+^ cells in thymus, skin and lymph nodes (Chen et al., 2019; Tan et al., 2019), how these subsets compare and how they are controlled in lung tissue is poorly understood.

Over the past few years, γδ T cells have garnered much attention in cancer, where unique subsets of γδ T cells can play either a pro-tumor or anti-tumor role (Silva-Santos et al., 2019). In mice, these functionally distinct γδ T cells can be stratified into two main subsets by the expression of the TNF receptor family member, CD27, which distinguishes IFNγ- from IL-17A-producing cells (Ribot et al., 2009). CD27^+^IFNγ^+^γδ T cells that express Vγ1 or Vγ4 TCR chains have anti-tumor properties capable of direct cancer cell killing and boosting CD8^+^ T cell cytotoxic responses (Chen et al., 2019; Dadi et al., 2016; Gao et al., 2003; He et al., 2010; Lanca et al., 2013; Liu et al., 2008; Riond et al., 2009; Street et al., 2004). By contrast, CD27^—^IL-17A^+^γδ T cells that express Vγ4 or Vγ6 TCR chains can drive primary tumor growth (Housseau et al., 2016; Jin et al., 2019; Ma et al., 2014; Patin et al., 2018; Rei et al., 2014; Rutkowski et al., 2015; Van Hede et al., 2017; Wakita et al., 2010), and they can promote metastasis (Baek et al., 2017; Benevides et al., 2015; Coffelt et al., 2015; Kulig et al., 2016; Wellenstein et al., 2019). One common mechanism that IL-17A- producing Vγ4^+^ and Vγ6^+^ cells share to foster cancer progression is the stimulation of granulopoiesis and recruitment of neutrophils to primary and secondary tumors that suppress anti-tumor CD8^+^ T cells. These immunosuppressive neutrophils are triggered by IL-17A- regulated G-CSF expression (Baek et al., 2017; Coffelt et al., 2015; Jin et al., 2019; Ma et al., 2014; Wellenstein et al., 2019). Within the lung microenvironment, γδ T cells are activated to produce IL-17A by tumor-derived or microbiota-induced IL-1β and IL-23 (Coffelt et al., 2015; Jin et al., 2019; Wellenstein et al., 2019), two cytokines with well-established influence on IL- 17A-producing γδ T cells (Sutton et al., 2009). In addition, these pathways are highly comparable to human cancer. IL-17A-producing γδ T cells not only infiltrate human tumors (Boufea et al., 2021; Kargl et al., 2017; Wu et al., 2014), but high levels of IL-17A or abundance of γδ T cells also correlates with poor prognosis and metastasis in cancer patients (Benevides et al., 2015; Ma et al., 2012; Wu et al., 2014). Collectively, these studies underpin the crucial role of lung IL-17A-producing γδ T cells in cancer progression. However, questions remain about how these cells are locally controlled in the tumor-conditioned lung microenvironment.

Here, we performed a comprehensive analysis of γδ T cell phenotype, transcriptional diversity and function in healthy lung and mammary tumor-conditioned lung (i.e. the pre-metastatic niche). Using single-cell RNA sequencing (scRNAseq) and flow cytometry, we show that Vγ6^+^ cells in healthy lung are distinguishable from Vγ4^+^ cells by expression of a number of molecules, including CXCR6, ICOS, JAML, NKG2D and PD-1. Manipulation of ICOS and PD-1 signaling on Vγ6^+^ cells affected IL-17A production as well as their proliferation. In a mouse model of breast cancer, scRNAseq revealed that the diversity of lung γδ T cells increases dramatically in response to a tumor. We found that both Vγ4^+^ and Vγ6^+^ cells proliferate and increase expression of IL-17A, IL-17F and AREG, which is mediated by IL-1β and IL-23. While ICOS and PD-1 expression remained at high levels on Vγ6^+^ cells in tumor-bearing mice, lung Vγ4^+^ cells up-regulated another co-inhibitory molecule, TIM-3. Furthermore, we demonstrate that inhibition of PD-1 or TIM-3, but not ICOS, enhances IL-17A-producing γδ T cells in the lung, indicating that PD-1 and TIM-3 function as inhibitory molecules on Vγ6^+^ cells and Vγ4^+^ cells, respectively. These data offer insight into the distinctive regulation of lung γδ T cell subsets in homeostasis and cancer.

## MATERIALS AND METHODS

### Mice

Female FVB/n mice (10-12 weeks old) were used throughout the study. These mice were bred at the CRUK Beatson Institute from the *K14-Cre;Brca1^F/F^;Trp53^F/F^* (KB1P) colony (Liu et al., 2007), which was gifted from Jos Jonkers (Netherlands Cancer Institute). KB1P mice were backcrossed onto the FVB/N background. *Cre* recombinase negative mice were used as wild-type (WT) sources for γδ T cell phenotyping and scRNA analysis. *Cre* recombinase positive mice were monitored twice weekly for tumor formation by palpation and calliper measurements starting at 4 months of age. Female C57BL/6J mice (10-12 weeks old) were also bred in house, and these mice were wild-type for every allele. For KB1P tumor transplantation studies, female FVB/n mice (6-8 weeks old) were purchased from Charles River (Cambridge, UK). Purchased mice were allowed to acclimatize before procedure until 10 weeks old. Transplantation of tumor fragments was performed as previously described (Millar et al., 2020). Once tumors reached 1 cm, mice were randomized into control or experimental groups. All animals were bred under specific pathogen-free conditions with unrestricted access to food and water. Procedures were performed in accordance with UK Home Office license numbers, 70/8645 and PP6345023 (to Karen Blyth), and they were carried out in line with the Animals (Scientific Procedures) Act 1986 and the EU Directive 2010 and sanctioned by Local Ethical Review Process (University of Glasgow).

### Single-cell RNA sequencing

Total γδ T cells were sorted from lungs of 4 WT FVB/n and 4 tumor-bearing KB1P mice by gating on DAPI^—^CD3^+^TCRδ^+^ cells (Supplemental Figure 1). Typically, 5000 γδ T cells were captured from 1 lung, using a BD FACSAria II Cell Sorter. The sorted cells were loaded onto the Chromium Single Cell 30 Chip Kit v2 (10xGenomics) to generate libraries for scRNAseq, following the manufacturer’s instructions. The sequencing-ready library was cleaned up with SPRIselect beads (Beckman Coulter, High Wycombe, UK). Quality control of the library was performed prior to sequencing (Qubit, Bioanalyzer, qPCR). Illumina sequencing was performed using NovaSeq S1 by Edinburgh Genomics (University of Edinburgh). The output .bcl2 file was converted to FASTQ format by using cellranger-mkfastq^TM^ algorithm (10xGenomics), and cellranger-count was used to align to the mm10 reference murine genome and build the final (cell, UMI) expression matrix for each sample. After quality control for removal of cells with over 3000 or less than 200 genes, and cells with more that 10% of reads from mitochondrial genes, followed by cell cycle correction, we obtained single cell transcriptomes of 3796 γδ T cells from lungs of WT mice and 5091 γδ T cells from lungs of mammary tumor-bearing KB1P mice. The data were normalised with the Seurat V3 “LogNormalize” method and batch effects where corrected with the Batchelor package. Then, principal component analysis (PCA) and unsupervised clustering were performed on the data by using Seurat. The first 20 principal components were retained for dimensional reduction and t-distributed Stochastic Neighbour Embedding (t-SNE) was utilized for visualization of the data. Marker genes and differentially expressed genes of cell clusters were determined by using Seurat and DESeq2, respectively. The signature scores were visualized as heatmap projected on the dataset t-SNE, with contours around those cells scoring > superior quartile. Raw data are deposited at ArrayExpress (E-MTAB-10677).

### Tissue processing

Lungs were mechanically dissociated using a scalpel and transferred to collagenase solution consisting of DMEM medium (ThermoFisher, Waltham, MA, USA) supplemented with 1 mg/mL collagenase D (Roche, Welwyn Garden City, UK) and 25 µg/mL DNase 1 (ThermoFisher). Enzymatic dissociation was assisted by heat and mechanical tissue disruption, using the gentleMACS Octo Dissociator, run: 37C_m_LDK_01 (Miltenyi Biotec, Surrey, UK), according to the manufacturer’s dissociation protocol. Tumors were enzymatically digested with Tumor Dissociation Kit, using the gentleMACS Octo Dissociator, run: 37C_m_TDK_01 (Miltenyi Biotec). The lung and tumor cell suspensions were filtered through a 70 µm cell strainer, using a syringe plunger, and enzyme activity was stopped by addition of 2 mL of fetal calf serum (FCS) followed by 5 mL DMEM medium supplemented with 10% FCS, 2 mM L-glutamine (ThermoFisher) and 10000 U/mL penicillin/streptomycin (ThermoFisher). Spleen and lymph nodes (LN) were forced through a 70 µm cell strainer, using a syringe plunger, and the tissue was flushed through with PBS containing 1% BSA.

Blood was drawn after cardiac puncture and collected in EDTA tubes and kept at room temperature for haematology analysis by Idexx (ProCyte Dx Haematology Analyser). For red blood cell lysis, the cell pellets were resuspended in 5 mL of commercially available 1x Red Blood Cell Lysis buffer (ThermoFisher) for 3 minutes. Cells were resuspended in PBS containing 0.5% BSA and cell number was acquired, using a haemocytometer.

### Flow cytometry

Single-cell suspensions were added to 96-well V bottom plates at a maximum density of 4×10^6^ cells/well. Cells were stimulated for 3 hours at 37 °C with complete IMDM medium with 1x Cell Activation Cocktail (with Brefeldin A, Biolegend). After stimulation, cells were centrifuged at 200g for 2 minutes. Cells were incubated in blocking buffer (50 µL PBS/0.5% BSA, 1 µL FcR block [Biolegend]) for 20 minutes at 4 °C. Antibodies for surface antigens were prepared in Brilliant stain buffer (BD Biosciences) and cells were stained for 30 minutes at 4 °C in the dark. Cells were washed with PBS/0.5% BSA, centrifuged at 200g for 2 minutes followed by ice-cold PBS and incubated with Zombie NIR Fixable Viability dye (Biolegend) to exclude dead cells for 20 minutes at 4 °C. After washing the cells with PBS/0.5% BSA, cells were fixed and permeabilized in FOXP3 Transcription Factor Fixation/Permeabilization solution (ThermoFisher) for 20 minutes at 4 °C, following the manufacturer’s instructions. Intracellular antibodies were prepared in permeabilization buffer and cells were incubated for 30 minutes at 4 °C. Cells were washed with permeabilization buffer, followed by PBS/0.5% BSA and resuspended in PBS/0.5% BSA. Fluorescence minus one (FMO) control samples were prepared from spare cells to discriminate positive and negative staining. All experiments were performed using a 5-laser BD LSRFortessa flow cytometer with DIVA software (BD Biosciences). Data were analyzed using FlowJo Software version 9.9.6. Antibodies for analysis are listed here:

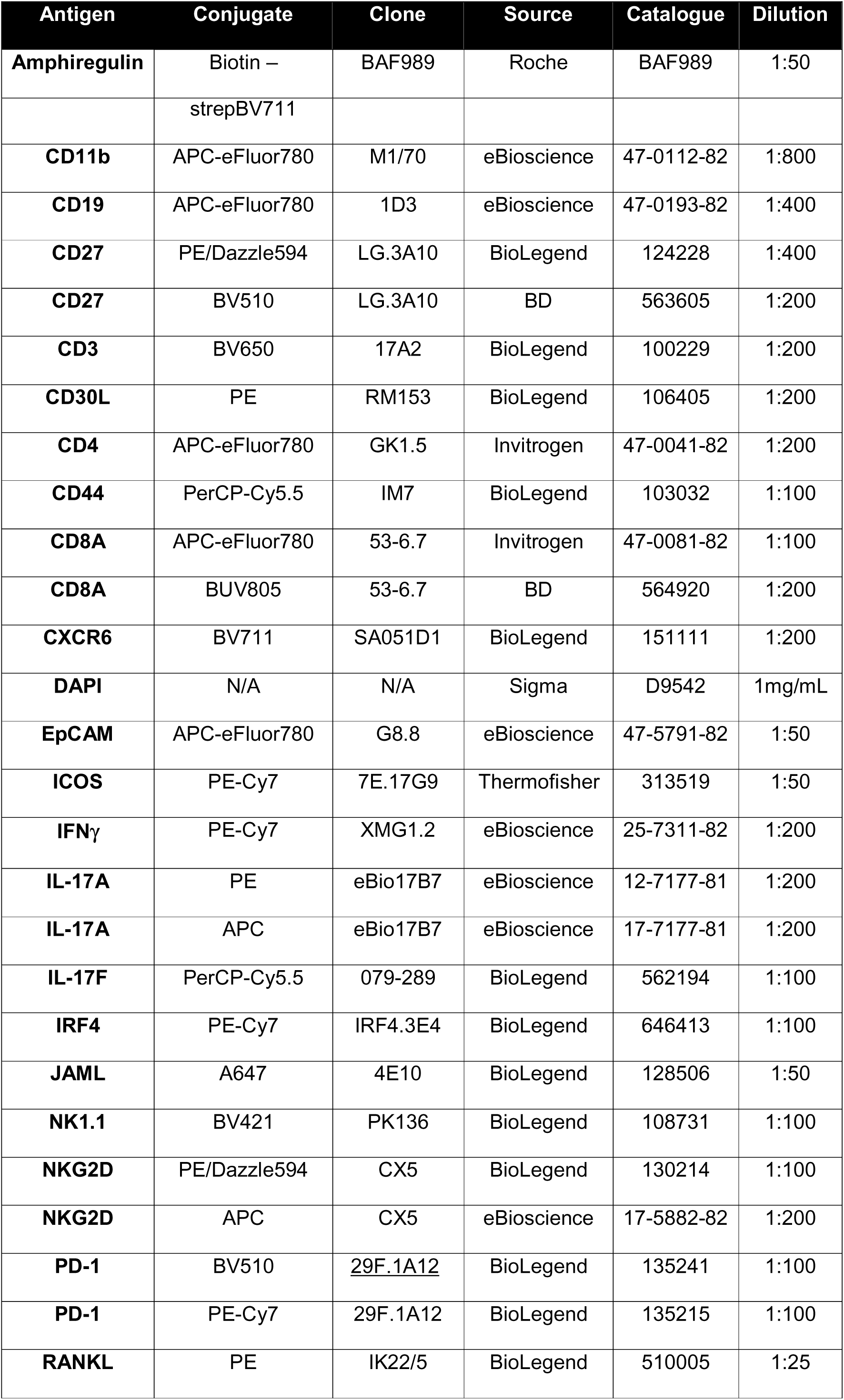
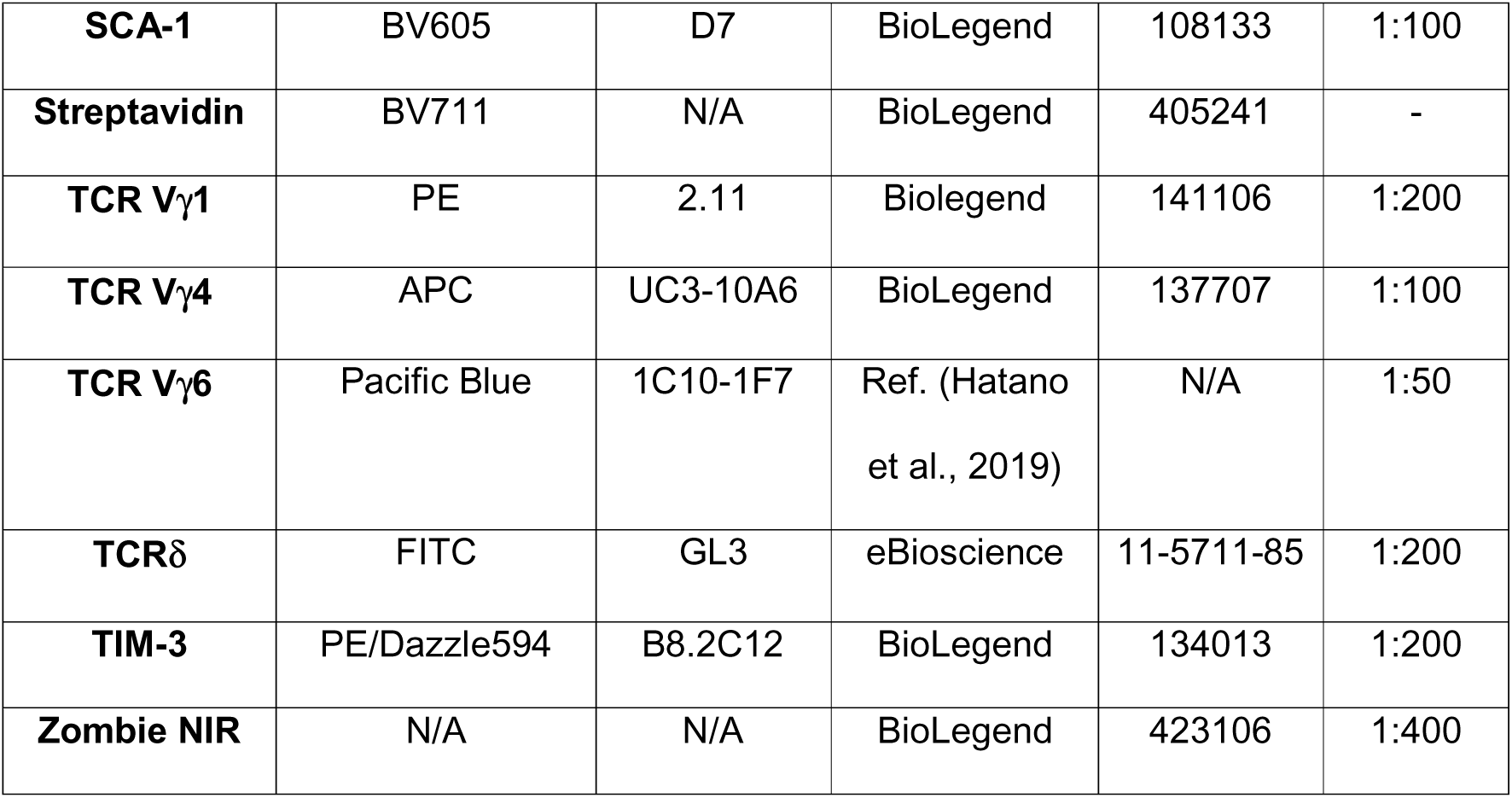

### *In vitro* T cell assays

CD3^+^ T cells from lung single-cell suspensions (1×10^7^ cells) were isolated by negative selection using the MojoSort mouse CD3^+^ T-cell Isolation kit (Biolegend) according to the manufacturer’s instructions. Cells after enrichment were counted with a haemocytometer and plated at a density of 1×10^6^ cells/mL in round bottom 96-well plates. CD3^+^ T cells were cultured for 15 hours or 24 hours as indicated. Cells were stimulated with 2.5 ng/mL or 5 ng/mL recombinant IL-1β (ImmunoTools, Friesoythe, Germany) and IL-23 (R&D Systems) in IMDM medium, 10% FCS, 2 mM L-glutamine, 10000 U/mL penicillin/streptomycin and 50 μM β-mercaptoethanol. Where indicated, wells were coated with 30 μg/mL PD-L1-Fc chimera protein (R&D Systems) or 6 μg/mL anti-ICOS (clone C398.4A, Biolegend) in PBS overnight at 4 °C. IL-17A levels in conditioned medium was determined by ELISA (R&D Systems) according to the manufacturer’s instructions. Each experiment was replicated at least 5 times.

### Blockade of PD-1, ICOS and TIM3 *in vivo*

Wild-type female FVB/n mice (Charles River), with or without transplanted KB1P tumors, were injected intraperitoneally with a single dose of 200 μg anti-PD-1 (clone RMP1-14, BioXcell), anti-ICOS (clone 7E.17G9, BioXcell), or anti-TIM-3 (clone RMT3-23, BioXcell), followed by single injections of 100 μg antibody on two consecutive days. Control mice followed the same dosage regime with isotype control (Rat IgG2a or Armenian Hamster IgG, BioXcell). Mice were sacrificed 24 hours after the third antibody injection.

### Statistical analysis

Non-parametric Mann-Whitney U test was used to compare two groups, while non-parametric Kruskal-Wallis followed by Dunn’s posthoc test was used to compare groups of three or more. Sample sizes for each experiment were based on a power calculation and/or previous experience of the mouse models. Analyses and data visualization were performed using GraphPad Prism (version 9.1.2) and Adobe Illustrator CS5.1 (version 15.1.0).

## RESULTS

### Single-cell RNA sequencing analysis identifies two clusters of γδ T cells in normal lung

Lung γδ T cells – particularly the IL-17A-producing cells – play important roles in homeostasis, infection, inflammation, allergy, cancer progression and metastasis (Coffelt et al., 2015; Faustino et al., 2020; Guo et al., 2018; Jin et al., 2019; Kulig et al., 2016; Misiak et al., 2017; Wang et al., 2021). However, knowledge on γδ T cell diversity, expression of distinguishing molecules and regulation within the lung is not well understood. To better understand γδ T cell heterogeneity in the lung, we performed single-cell RNA sequencing (scRNAseq) of total CD3^+^TCRδ^+^ cells isolated from the lungs of wild-type (WT) FVB/n mice (**Supplemental Figure 1**). The isolated fraction comprised 2.5-5.3% of the viable cells. Libraries for scRNAseq were prepared using the Chromium 10X platform and we obtained single-cell transcriptomes of 3796 γδ T cells from lungs of WT mice. Principal component analysis (PCA) for dimensional reduction and unsupervised clustering were performed on the data, and t-Distributed Stochastic Neighbour Embedding (tSNE) was utilized for visualization of the data in two dimensions (**Figure 1A**). This unbiased transcriptional analysis of 3796 individual cells segregated γδ T cells into two major clusters, which were designated as Cluster 1 and Cluster 2. Cluster 1 represented the largest number of γδ T cells within the lung compartment. Cluster 2 consisted of two transcriptionally related, yet distinct groups, which we labelled Cluster 2.1 and 2.2 **(Figure 1A)**. Expression of *Cd3d* and *Cd3e* genes confirmed that all of these cells are TCR-expressing cells **(Figure 1B)**. To identify distinguishing characteristics separating Cluster 1 and Cluster 2, expression of a selected set of established marker genes often used to discriminate IL-17A-producing γδ T cells from IFNγ- producing γδ T cells was investigated within the scRNAseq dataset. CD27, a co-stimulatory molecule, was chosen because its protein expression on γδ T cells stratifies IL-17A- producing (CD27^—^) cells from IFNγ-producing (CD27^+^) cells (Ribot et al., 2009). Expression of *Cd27* was largely absent from Cluster 1 and enriched in Clusters 2.1 and 2.2 (**Figure 1C**), suggesting that Cluster 1 may represent IL-17A-producing cells while Cluster 2 may represent IFNγ-producing cells. This hypothesis was confirmed when enrichment of *Il17a* and other genes associated with IL-17A signaling was observed in Cluster 1, such as the cytokine receptors, *Il23r* and *Il1r1*, as well as the transcription factors, *Rorc* and *Maf* **(Figure 1C)**. Expression of *Ifng* and the gene encoding the transcription factor that regulates *Ifng* expression, *Tbx21* (T-Bet), were both localized to Cluster 2.2 and dispersed throughout Cluster 1 **(Supplemental Figure 2A**). Expression of *Cd28*, which encodes a co-stimulatory molecule involved in IFNγ-producing γδ T cell expansion and IL-2 production (Ribot et al., 2012), was enriched in Cluster 2 **(Supplemental Figure 2A**). These data indicate that the two major clusters of lung γδ T cells identified by scRNAseq are largely defined by *Cd27* expression and IL-17A signaling molecules in accordance with historical literature (Ciofani et al., 2012; Petermann et al., 2010; Ribot et al., 2009; Sutton et al., 2009; Zuberbuehler et al., 2019).

**Figure 1:**
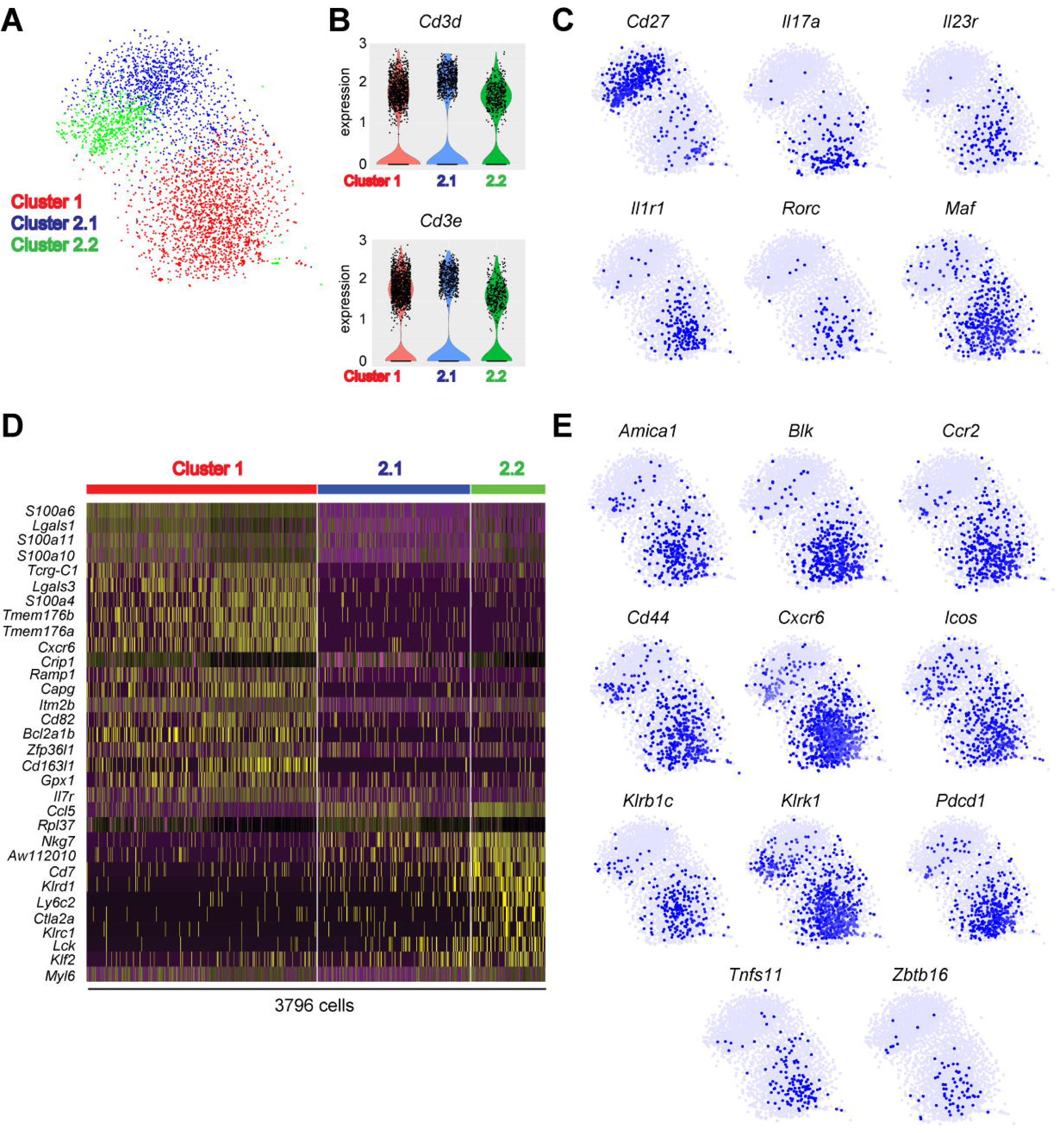
Single-cell RNA sequencing identifies two major clusters of lung. γδ **T cells. (A)** Two-dimensional visualization of single γδ T cells from lungs of three wild-type FVB/n mice via tSNE. Each dot represents an individual cell. **(B)** Violin plots of *Cd3d* and *Cd3e* expression for Clusters 1, 2.1 and 2.2. **(C)** Feature plots in tSNE map of indicated genes. **(D)** Heatmap showing z-score normalized expression of top 32 differentially expressed genes for Clusters 1, 2.1 and 2.2. Cells are plotted in columns by cluster, and genes are shown in rows. Gene expression is colour coded with a scale based on z-score distribution, from -2 (purple) to 2 (yellow). **(E)** Feature plots in tSNE map of indicated genes.

To uncover additional transcriptional differences between Clusters 1 and 2, differentially expressed genes defining each cluster were analyzed. The top 50 most significant genes from each cluster were combined to generate a heat map consisting of 32 genes (**Figure 1D; Table 1**). The clusters were defined by diverse expression profiles for TCR signaling, cytokines, cytokine receptors, chemokine receptors, co-stimulatory and co-inhibitory molecules as well as NK cell markers **(Figure 1D)**. Cluster 1 representing CD27^—^ γδ T cells was enriched in genes from the S100 family of calcium-binding proteins, including *S100a4, S100a6, S100a10* and *S100a11* (**Figure 1D**). Members of the Galectin family, including *Lgals1* (Galectin-1) and *Lgals3* (Galectin-3), which are carbohydrate binding proteins, were enriched in Cluster 1 (**Figure 1D**). Galectin-1-expressing γδ T cells are known to suppress antigen-specific anti-tumor immunity in a Toll-like receptor 5 (TLR5)-dependent manner (Rutkowski et al., 2015). The ion channel homologues, *Tmem176a* and *Tmem176b*, were defining genes of Cluster 1 (**Figure 1D**). Expression of *Tmem176a* and *Tmem176b* is regulated by the *Il17a*-inducing transcription factor, RORγt (Ciofani et al., 2012). TMEM176A and TMEM176B have redundant functions in controlling ion influx/efflux in γδ T cells (Drujont et al., 2016). The chemokine receptor, *Cxcr6*, and the cytokine receptor, *Il7r*, appeared in the top 50 list of genes. Both receptors are established regulators of IL-17A-producing γδ T cell trafficking and cytokine expression (Butcher et al., 2016; Chen et al., 2019; Gray et al., 2012; Gray et al., 2011; Hayes et al., 1996; Michel et al., 2012). Cells in Cluster 1 expressed *Bcl2a1b* as well as *Bcl2a1a* and *Bcl2a1d* (**Table 1**), which are pro-survival factors that support Vγ6^+^ cells in skin (Tan et al., 2019). The other top genes for Cluster 1 included *Crip1, Ramp1, Capg, Itm2b, Cd82, Zfp36l1,* and *Gpx1* (**Figure 1D**). Vγ6^+^ cells makeup the largest proportion of γδ T cells in the lung (Hayes et al., 1996; McIntyre et al., 2020). In agreement with these data, markers of Vγ6^+^ cells appeared in Cluster 1, which was the cluster containing the greatest number of cells (**Figure 1A**). These markers included *Tcrg-C1* and *Cd163l1,* which encodes the SCART1 protein (**Figure 1D, Supplemental Figure 2B**) (Chen et al., 2019; Kisielow et al., 2008; Tan et al., 2019). The markers of Vγ4^+^ cells, including *5830411N06Rik* (the gene encoding SCART2), *Sox13* and *Cd9* (Chen et al., 2019; Gray et al., 2013; Malhotra et al., 2013; Tan et al., 2019), were expressed by only a few cells in Cluster 1 (**Supplemental Figure 2B**). Together, the defining genes of Cluster 1 identified for lung γδ T cells (e.g. *Tcrg-C1, Cd163l1, Cd27, Maf, S100a* genes*, Lgals1 and Lgals1, Bcl2a1* genes, *Cxcr6,* etc.) are highly similar to the transcriptome of IL-17A-producing Vγ6^+^ cells from skin, thymus, adipose, uterus and lymph nodes (Chen et al., 2019; Kohlgruber et al., 2018; Monin et al., 2020; Tan et al., 2019), suggesting that these commonalities can be used to identify Vγ6^+^ cells across various tissues.

Further investigation into the characteristics of Cluster 1 revealed additional insight into the transcriptome of these mostly Vγ6^+^ cells. Common genes for IL-17A-producing γδ T cells, such as *Blk, Cd44, Ccr2,* and *Zbtb16* (which encodes PLZF) (Ciofani et al., 2012; Kohlgruber et al., 2018; Laird et al., 2010; McKenzie et al., 2017; Stark et al., 2005), were readily expressed in Cluster 1 cells (**Figure 1E**). Expression of the chemokine receptor, *Cxcr6*, and *Tnfsf11*, which encodes the cytokine, RANKL, was more prevalent in Cluster 1 than Cluster 2 (**Figure 1E**). Two NK cell-associated molecules, *Klrb1c* and *Klrk1*, were highly enriched in Cluster 1. This observation was unexpected because these genes encode NK1.1/ CD161 and NKG2D, respectively, which are two proteins known for their involvement in target recognition and cancer cell killing. In addition, *Amica1* and *Icos*, two co-stimulatory receptors, and *Pdcd1* (PD-1), a co-inhibitory receptor, were enriched in Cluster 1 (**Figure 1E**). The *Amica1* gene encodes JAML, which is an activator of skin-resident, Vγ5 cells (Witherden et al., 2010). ICOS signaling is important in thymic development of IL-17A- producing γδ T cells and function during experimental autoimmune encephalomyelitis (EAE) (Buus et al., 2016; Galicia et al., 2009), while the function of PD-1 on these cells is unknown. Apart from the NK cell-associated genes, the Cluster 1 phenotype (e.g. *Amica1*, *Cxcr6, Icos, Pdcd1* and *Tnfsf11*) was highly reminiscent of tissue resident memory (T_rm_) T cells (Clarke et al., 2019; Djenidi et al., 2015; Kumar et al., 2017; Mackay et al., 2013; Strutt et al., 2018; Wein et al., 2019), indicating that lung resident Vγ6^+^ cells share many commonalities with antigen-experienced, non-circulating αβ T cells in the lung.

The defining genes of Cluster 2 represented TCR signaling molecules, cytotoxic molecules, cytokines, and T cell effector molecules, consistent with their CD27-expressing status and the established function of IFNγ-producing γδ T cells (Silva-Santos et al., 2019). Some of these genes included *Ccl5, Rpl37, Nkg7, AW112010, Cd7, Klrd1, Ly6c2, Ctla2a, Klrc1, Lck, Klf2* and *Myl6* (**Figure 1D**). The product of the *Ly6c2* gene, Ly6C, is a molecule more commonly associated with monocytes and neutrophils, and it can be expressed by CD27^+^ γδ T cells (Lombes et al., 2015); although, its function on these cells is unknown. *Ly6c2* was expressed by Cluster 2.2 (**Supplemental Figure 2C**). The KLF2 transcription factor is known to regulate γδ T cell trafficking via expression of sphingosine 1-phosphate receptor 1 (S1PR1) (Odumade et al., 2010; Ugur et al., 2018). In addition to the top 50 genes (**Figure 1D**), the migratory-related molecules, *Ccr7* and *S1pr1*, were also a predominant feature of Cluster 2 with specific enrichment in Cluster 2.1 (**Supplemental Figure 2C**). Taken together, these data uncover novel heterogeneity within the CD27^+^ compartment of γδ T cells, and they suggest that cells in Cluster 2.2 may represent a more activated effector population of CD27^+^ γδ T cells.

### Lung CD27^—^ γδ T cells display a tissue resident memory phenotype

Having identified major transcriptional differences between lung γδ T cells by scRNAseq, we validated some of these differences at the protein level. We focused on the observation that cells from Cluster 1 expressed genes associated with T_rm_ cells (e.g. *Amica1*, *Cxcr6, Icos, Pdcd1* and *Tnfsf11*) as well as the NK cell markers, *Klrb1c* (NK1.1/CD161) and *Klrk1* (NKG2D). Flow cytometry analysis of lung γδ T cells isolated from FVB/n mice revealed that CD27^—^ γδ T cells expressed higher levels of CXCR6, ICOS, JAML, PD-1 and RANKL, when compared to lung CD27^+^ γδ T cells **(Figure 2A, B)**, confirming the scRNAseq data and the similarity with T_rm_ cells (Clarke et al., 2019; Djenidi et al., 2015; Kumar et al., 2017; Mackay et al., 2013; Strutt et al., 2018; Wein et al., 2019). The NK cell-associated killing receptors, NK1.1 and NKG2D, were also expressed more abundantly on CD27^—^ γδ T cells compared with CD27^+^ γδ T cells **(Figure 2A, B)**. These data suggest that NKG2D and NK1.1 signaling may regulate lung CD27^—^ γδ T cells. In support of this notion, blockade of NKG2D reduces IL-17A expression by tumor-infiltrating γδ T cells (Tang et al., 2013; Wakita et al., 2010).

**Figure 2:**
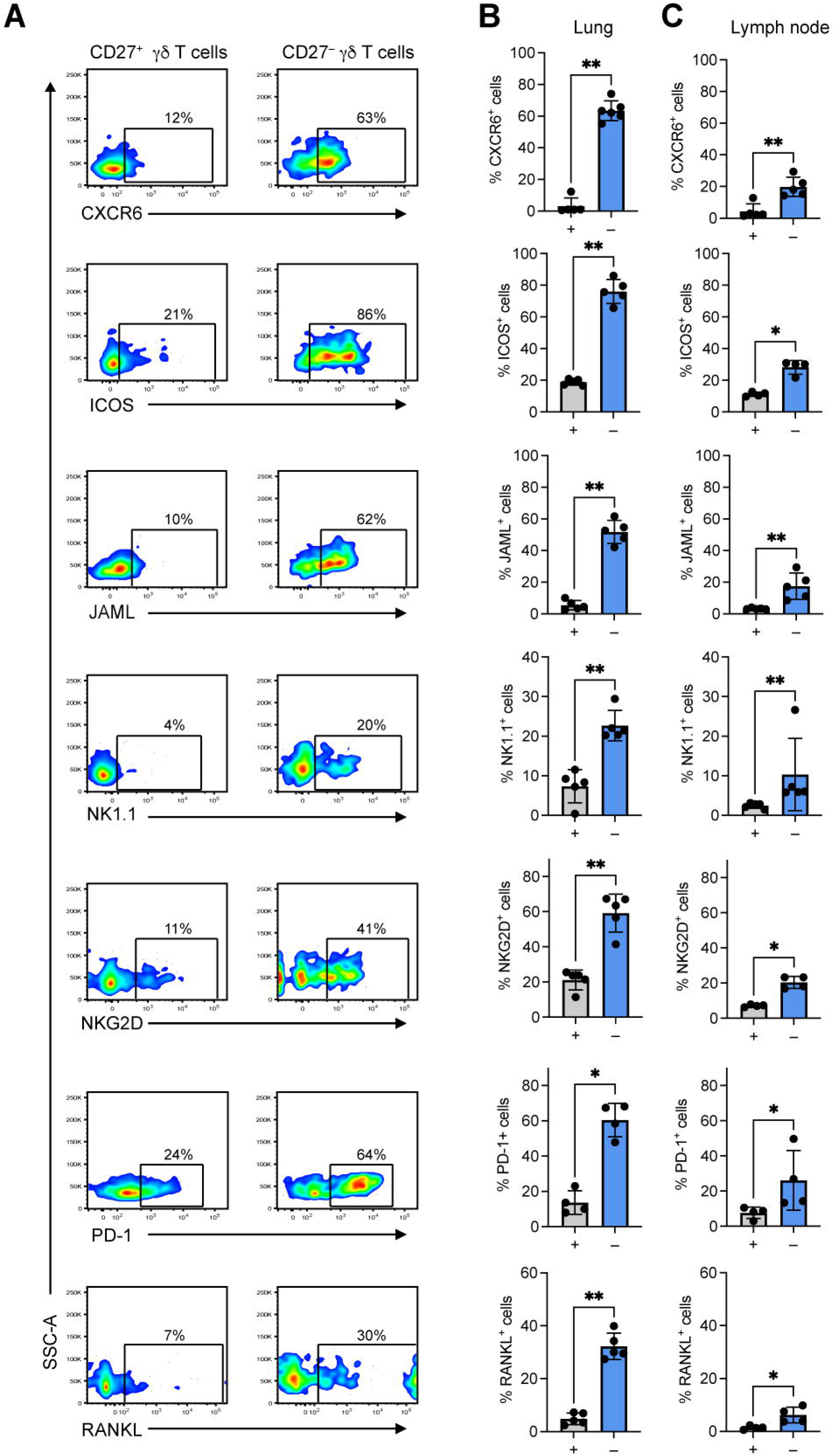
Lung CD27^—^ γδ T cells express a tissue-resident memory phenotype. Single cell suspensions of lung and lymph nodes from FVB/n mice were stained with antibodies against CD3, TCRδ, CD27 and the indicated molecules. Cells were analyzed by flow cytometry. **(A)** Representative flow cytometry plots of indicated molecule expression in CD27^+^ and CD27^—^ γδ T cells. Numbers indicate percentage positive cells in gate. **(B, C)** Graphic representation of percentage positive cells for each indicated molecule in CD27^+^ (“+”) and CD27^—^ (“—“) γδ T cells from lung and lymph node. Each dot represents one mouse. Data are presented as mean ± SD. \**p* < 0.05, \*\**p* < 0.01 as determined by Mann-Whitney U test.

We also compared expression of these T_rm_ markers on CD27^—^ and CD27^+^ γδ T cells from the lymph node, which contains higher numbers of CD27^—^Vγ4^+^ cells and lower numbers of CD27^—^Vγ6^+^ cells than the lung. This analysis revealed that CXCR6, ICOS, JAML, NK1.1, NKG2D, PD-1 and RANKL are more highly expressed on CD27^—^ γδ T cells in a similar manner to cells from lung (**Figure 2C**). The reduced level of expression may reflect lower numbers of CD27^—^Vγ6^+^ cells in lymph nodes.

NK1.1/CD161 is an established marker of IFNγ-producing CD27^+^ γδ T cells that distinguishes these cells from IL-17A-producing γδ T cells (Haas et al., 2009). Therefore, we hypothesized that the discrepancy between our data from lung and established literature may be explained by the differences in mouse strains. To this end, we analyzed lung γδ T cells in C57BL/6J mice for the same proteins. Lung CD27^—^ γδ T cells from C57BL/6J mice expressed higher levels of CXCR6, ICOS, JAML, NKG2D and RANKL, when compared to lung CD27^+^ γδ T cells (**Supplemental Figure 3**), corroborating the observations in FVB/n mice. In contrast to FVB/n mice, however, NK1.1 expression was lower on CD27^—^ γδ T cells than CD27^+^ γδ T cells from C57BL/6J mice **(Supplemental Figure 3**). These data indicate that the T_rm_ phenotype of CD27^—^ γδ T cells is stable between mouse strains, while expression of NK1.1 (and likely other molecules) is dissimilar between strains.

### PD-1 regulates IL-17A expression in lung Vγ6^+^ T cells

Given the high expression of ICOS and PD-1 on CD27^—^ γδ T cells (Figures 1 and 2), we examined whether expression of these molecules was specific to a particular γδ T cell subset. Flow cytometry analysis of lung CD27^—^ γδ T cells from WT mice revealed that ICOS and PD-1 are primarily expressed by Vγ6^+^ cells, whereas only a minor proportion of Vγ4^+^ cells expressed ICOS and PD-1 (**Figure 3A, B**). These data indicate that Vγ6^+^ cells produce constitutive levels of ICOS and PD-1 in steady state. We compared the phenotype of Vγ6^+^ICOS^+^PD-1^+^CD27^—^ cells versus Vγ4^+^ICOS^—^PD-1^—^CD27^—^ cells to understand how these cells differ in homeostatic lung tissue. Vγ6^+^ICOS^+^PD-1^+^CD27^—^ cells expressed higher levels of CXCR6, IL-17A, JAML, NK1.1 and NKG2D than Vγ4^+^ICOS^—^PD-1^—^CD27^—^ cells (**Figure 3C**). These data suggest that Vγ6^+^ cells and Vγ4^+^ cells are governed by different molecules in homeostatic lung.

**Figure 3:**
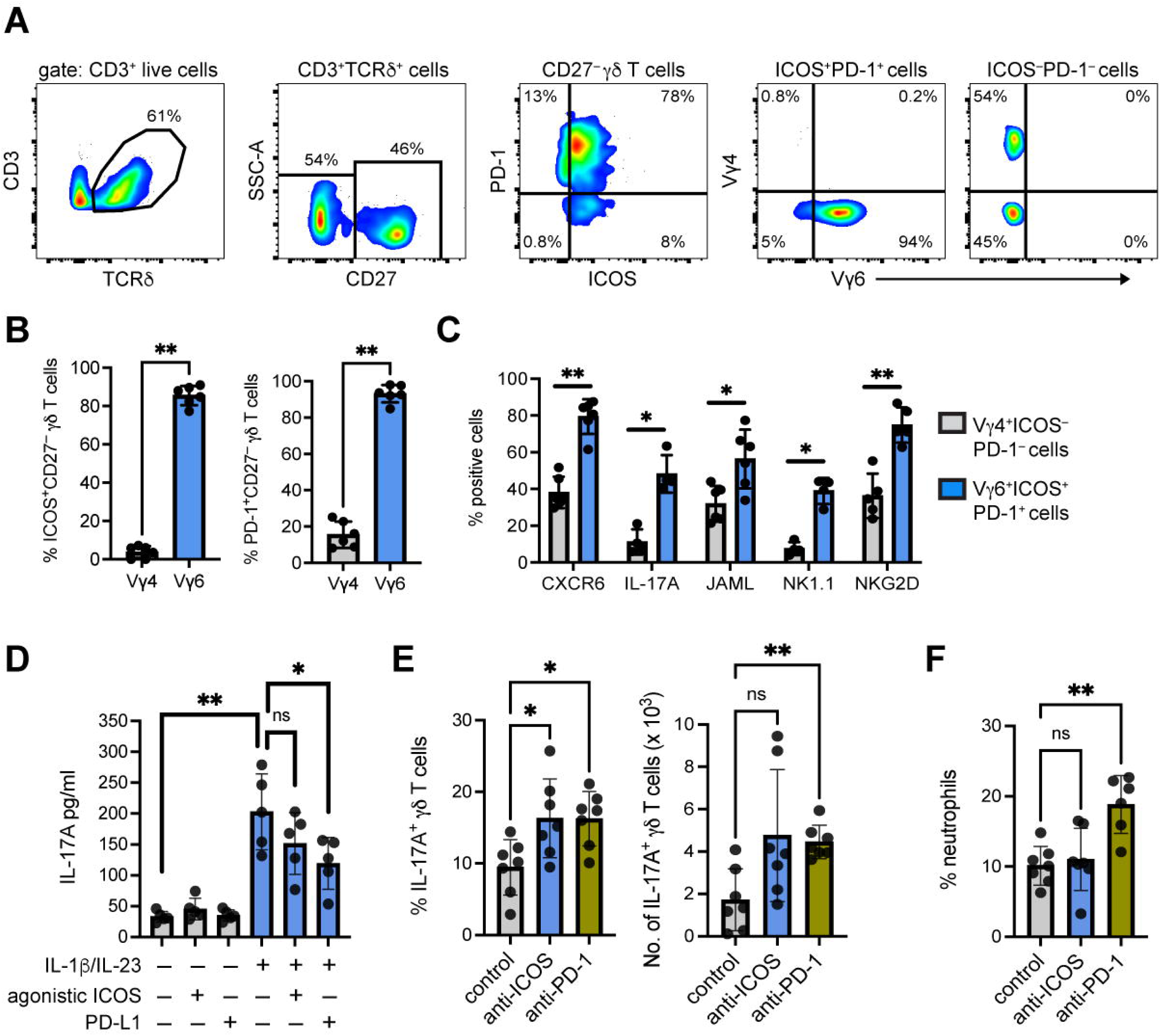
PD-1 signaling regulates IL-17A expression in lung Vγ6^+^ cells. **(A)** Gating strategy for ICOS and PD-1 expression on lung γδ T cell subsets from FVB/n mice. **(B)** Percentage of PD-1 and ICOS expression by lung Vγ1^+^, Vγ4^+^ and Vγ6^+^ CD27^—^ cells. **(C)** Expression of CXCR6, JAML, IL-17A, NK1.1 and NKG2D by Vγ4^+^ICOS^—^PD-1^—^ CD27^—^ cells (gray) and Vγ6^+^ICOS^+^PD-1^+^CD27^—^ cells (blue). **(D)** CD3^+^ T cells were isolated from the lungs of FVB/n mice and stimulated with recombinant IL-1β and IL-23 in the presence of plate-bound PD-L1-Fc or plate-bound anti-ICOS for 24 hours. Supernatants were examined for IL-17A levels by ELISA. **(E)** Wild-type FVB/n mice were injected with a single dose of 200 μg anti-PD-1 or anti-ICOS followed by injections of 100 μg for two consecutive days. Control mice followed the same dosage regime with isotype control. Mice were sacrificed 24 hours after the third injection. Single cell suspensions from lung were stimulated for 3 hours with PMA, ionomycin and Brefeldin A. Cells were stained with antibodies against CD3, TCRδ, CD27 and IL-17A to identify CD27^—^ γδ T cells. The proportion of cells expressing IL-17A and the absolute number of cells is represented graphically. **(F)** Percentage of neutrophils in isotype control, anti-PD-1- and anti-ICOS-treated mice as measured by IDEXX ProCyte hematology analyzer. Each dot represents one mouse. Data are presented as mean ± SD. \**p* < 0.05, \*\**p* < 0.01 as determined by Mann-Whitney U test or Kruskal-Wallis test followed by Dunn’s posthoc test.

PD-1, through its interaction with PD-L1 or PD-L2, functions as a negative regulator of T cell activation by interfering with TCR signaling (Honda et al., 2014). ICOS is a co-stimulatory molecule primarily expressed by CD4^+^ T helper and regulatory cells (Amatore et al., 2020), but it is also important for development and regulation of IL-17A-producing γδ T cells (Buus et al., 2016; Galicia et al., 2009). Therefore, we hypothesized that ICOS and PD- 1 are negative regulators of Vγ6^+^ cells. To address this hypothesis, we stimulated lung CD3^+^ T cells with IL-1β and IL-23 in the presence of recombinant PD-L1 or an ICOS agonistic antibody and measured IL-17A by ELISA. IL-17A levels in conditioned medium were increased by IL-1β and IL-23 stimulation. However, the addition of PD-L1 co-stimulation diminished IL-1β/IL-23-induced IL-17A production, while ICOS co-stimulation failed to significantly affect IL-17A production **(Figure 3D)**. These data indicate that activation of the PD-1 pathway functions to limit IL-17A expression in lung γδ T cells. Consequently, we examined the effect of anti-PD-1 and anti-ICOS blocking antibodies on lung γδ T cells *in vivo*. Naïve FVB/n mice were treated for three consecutive days with anti-PD-1, anti-ICOS or isotype control antibody, then we measured IL-17A expression in lung γδ T cells by intracellular flow cytometry. In this experiment, inhibition of PD-1 increased the frequency and absolute number of lung IL-17A-producing γδ T cells, whereas inhibition of ICOS increased the frequency but not the absolute number of lung IL-17A-producing γδ T cells **(Figure 3E)**. Since IL-17A is an essential cytokine for neutrophil expansion (Coffelt et al., 2015), we measured neutrophils in the blood of isotype control-, anti-PD-1- and anti-ICOS- treated WT mice. We found that the frequency of blood neutrophils is increased by anti-PD-1 treatment but anti-ICOS treatment failed to affect neutrophils when compared to controls **(Figure 3F)**. Taken together, these data indicate that PD-1 is a negative regulator of lung Vγ6^+^ cells, whose activation limits IL-17A expression in these cells, while the role of ICOS signaling on these cells remains unclear.

### The tissue resident memory phenotype of lung CD27^—^ γδ T cells is largely unaffected by tumor-derived factors

Since lung γδ T cells are important mediators of cancer progression and metastasis (Coffelt et al., 2015; Jin et al., 2019; Kulig et al., 2016), we profiled these cells for expression of ICOS and PD-1 as well as the other molecules identified in scRNAseq using a mouse model of breast cancer in which the γδ T cell – IL-17A – neutrophil axis is established (Wellenstein et al., 2019). We used the *K14-Cre;Brca1^F/F^;Trp53^F/F^* (KB1P) model of triple negative breast cancer, which develops invasive ductal carcinoma in one or more mammary glands at around eight months of age (Liu et al., 2007). When comparing tumor-bearing KB1P mice with tumor-free WT mice, the proportion of CD27^—^ γδ T cells and the total number of Vγ1^+^, Vγ4^+^ and Vγ6^+^ cells was higher in the lungs of tumor-bearing mice (**Figure 4A, B**). These data indicate that tumors in the mammary gland have influence over γδ T cells in the pre-metastatic lung. Further analysis revealed that mammary tumors increase ICOS expression and decrease NK1.1 expression on CD27^—^ γδ T cells, while expression of CXCR6, JAML, NKG2D and PD-1 remain consistent with WT mice (**Figure 4C**). These data suggest that the T_rm_ markers identified for lung Vγ6^+^ cells are largely constant between WT and mammary tumor-bearing mice.

**Figure 4:**
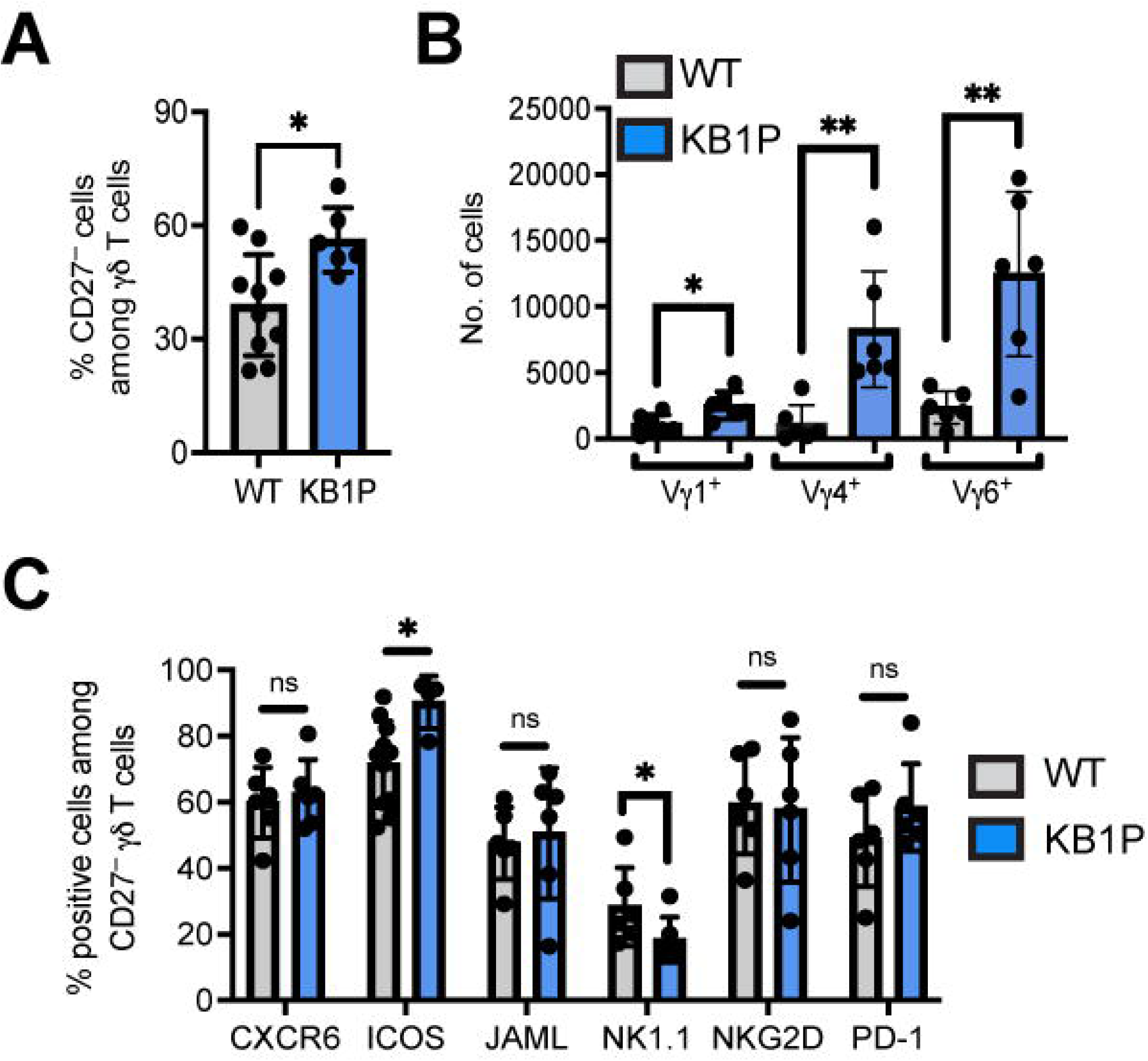
Lung γδ T cells expand in mammary tumor-bearing mice, but their tissue resident phenotype remains stable. **(A)** Frequency of CD27^—^ γδ T cells in wild-type (WT) and tumor-bearing *K14- Cre;Brca1^F/F^;Trp53^F/F^* (KB1P) mice. **(B)** Absolute number of Vγ1^+^, Vγ4^+^ and Vγ6^+^ CD27^—^ cells in the lungs of WT and KB1P mice. **(C)** Expression of CXCR6, ICOS, JAML, NK1.1, NKG2D and PD-1 by lung CD27^—^ γδ T cells from WT and tumor-bearing KB1P mice. Each dot represents one mouse. Data are presented as mean ± SD. \**p* < 0.05, \*\**p* < 0.01 as determined by Mann-Whitney U test.

### Lung γδ T cell diversity increases in response to mammary tumors

To gain a deeper understanding of lung Vγ4^+^ and Vγ6^+^ cell phenotype in KB1P tumor-bearing mice, we performed scRNAseq analysis on total lung γδ T cells from KB1P mice. From the computational interrogation of 5091 individual cells, clustering analysis revealed that lung γδ T cells segregate into 7 unique clusters (**Figure 5A**), which was remarkably different from the 2 clusters observed in WT mice (**Figure 1A**). Clusters 1-6 mostly lacked expression of *Cd27* mRNA, whereas Cluster 7 was enriched in cells expressing *Cd27* (**Figure 5B**), indicating that lung CD27^—^ γδ T cells are highly responsive to mammary tumors. This observation reflected the expansion of CD27^—^ γδ T cells we found by flow cytometry (**Figure 4A**). We evaluated the data for expression of genes identified from WT scRNAseq analysis and other common marker genes for IL-17A-producing γδ T cells, including *Amica1, Blk, Ccr2, Cd44, Cxcr6, Icos, Il1r1, Il23r, Klrbc1, Klrk1, Maf, Pdcd1, Rorc* and *Tnfsf11*. This investigation showed that these genes are primarily localized to Clusters 1-6 (**Figure 5B**), reinforcing the notion that Clusters 1-6 represent CD27^—^ γδ T cells.

**Figure 5:**
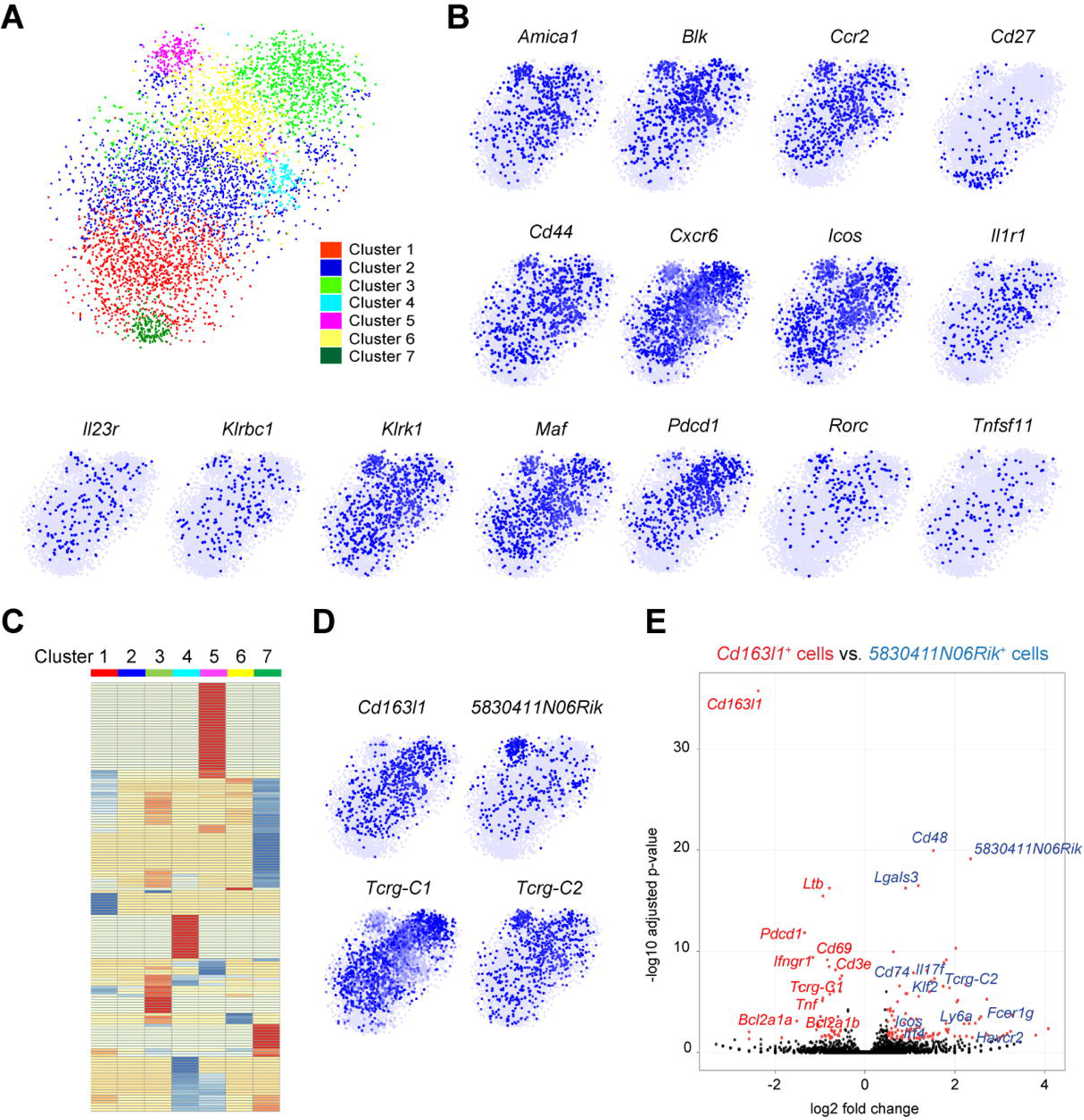
Mammary tumors instigate increased heterogeneity of lung γδ T cells. **(A)** Two-dimensional visualization of single γδ T cells from lungs of three tumor-bearing KB1P mice via tSNE. **(B)** Feature plots in tSNE map of indicated genes. **(C)** Heatmap showing z-score normalized expression of 278 differentially expressed genes for Clusters 1 to 7 (from Figure 4C). Clusters are plotted in columns, and genes are shown in rows. Gene expression is color coded with a scale based on z-score distribution, from -2 (blue) to 2 (red). **(D)** Feature plots in tSNE map of indicated genes. **(E)** The transcriptome of *Cd163l1*- and *5830411N06Rik*-expressing cells from 5A was compared with each other and represented as a volcano plot. Genes highly expressed by *Cd163l1*-expressing cells are denoted in red, and genes highly expressed by *5830411N06Rik*-expressing cells are denoted in blue.

Having observed a dramatic increase in transcriptional diversity among lung γδ T cells from tumor-bearing KB1P mice, we investigated the gene expression differences between individual clusters. Of the 593 differentially expressed genes highlighted in the clustering analysis, 497 genes (84%) were shared between 2 or more clusters. We noticed that 315 of these shared genes were ribosomal-related genes (**Table 2**). After excluding these ribosome genes, we generated a heat-map of the remaining 278 genes (**Figure 5C**) and we found that 184 genes (74%) were shared between 2 or more clusters. Clusters 3, 4, 5 and 7 were the most distinctive groups of cells, whereas Clusters 1, 2 and 6 were very similar to other clusters (**Figure 5C**). The genes unique to each individual cluster included 8 genes for Cluster 1, 0 genes for Cluster 2, 7 genes for Cluster 3, 22 genes for Cluster 4, 34 genes for Cluster 5, 2 genes for Cluster 6 and 21 genes for Cluster 7.

Both shared and unique genes for each cluster were analyzed in REACTOME pathway analysis to gain a better understanding of how cluster-specific genes are related. Cluster 1 was defined by autophagy-related pathways. Clusters 2 and 3 were defined by IL-12 and JAK/STAT signaling pathways. Cluster 4 was defined by interferon signaling and neutrophil degranulation. Cluster 5 was defined by IL-4/IL-13 signaling, RUNX2 gene regulation, platelet degranulation as well as iron uptake and transport. Cluster 6 was defined by RUNX1 gene regulation and FLT3/STAT5 signaling. The CD27^+^ γδ T cell cluster 7 was defined by cell responses to chemical stress, viruses or reactive oxygen species (**Supplemental Figure 4; Table 3**). Taken together, the data show that each cluster is transcriptionally distinct from the others, but overall, the clusters are very similar to each other, especially for Clusters 1-6.

Since Cluster 5 displayed the most unique set of genes, we determined what was driving these distinctive properties. Investigation of the Cluster 5 gene list revealed that *5830411N06Rik* (SCART2) and *Tcrg-C2,* two surrogate markers of Vγ4^+^ cells (Chen et al., 2019; Kisielow et al., 2008; Tan et al., 2019), are highly expressed (**Figure 5D**). We then used *Cd163l1* (SCART1) and *Tcrg-C1* to identify Vγ6^+^ cells, which showed that these cells are enriched in Clusters 1, 2, 3, 4 and 6, making up the majority of cells in the scRNAseq dataset. Vγ4^+^ and Vγ6^+^ cell markers were mostly absent from Cluster 7 **(Figure 5D)**. To compare the transcriptional profile of Vγ4^+^ and Vγ6^+^ cells from lungs of KB1P tumor-bearing mice, *5830411N06Rik* and *Cd163l1* genes were used to distinguish the two subsets. Computational analysis revealed that there were 106 genes unique to *5830411N06Rik*- expressing and 46 genes unique to *Cd163l1*-expressing cells (**Table 4**). Among the differentially expressed genes, *5830411N06Rik*-expressing cells produced higher levels of *Cd48, Cd74, Fcer1g, Havcr2, Icos, Il17f*, *Irf4, Klf2*, *Lgals3, Ly6a* and *Tcrg-C2,* while *Cd163l1*- expressing cells produced higher levels of *Bcl2a1a, Bcl2a1b, Cd3e, Cd69, Ifngr1, Ltb, Pdcd1, Tcrg-C1* and *Tnf.* Moreover, *Il17a* was equally expressed between the two subsets (**Figure 5E; Table 4**). The gene signatures of the two γδ T cell subsets mostly corroborate observations made in skin Vγ4^+^ and Vγ6^+^ cells; although, there were some exceptions, such as the enrichment of *Ifngr1* in skin *5830411N06Rik*-expressing cells and lung *Cd163l1*- expressing cells (Tan et al., 2019). These dissimilarities between lung and skin cells may reflect the influence of tumor-derived factors on lung Vγ4^+^ and Vγ6^+^ cells. Indeed, lung Vγ4^+^ and Vγ6^+^ cells from tumor-bearing KB1P mice showed expression of *Havcr2, Icos*, *Ly6a* and *Irf4*, for example (**Figure 5E**), which was not observed in cells from WT mice (Figure 1).

### IL-1β and IL-23 drive the phenotypic diversity of lung γδ T cells

Several studies in mouse models of cancer show that CD27^—^ γδ T cells upregulate IL-17A in response to tumor-derived factors (Baek et al., 2017; Benevides et al., 2015; Coffelt et al., 2015; Housseau et al., 2016; Jin et al., 2019; Kulig et al., 2016; Ma et al., 2014; Patin et al., 2018; Rei et al., 2014; Rutkowski et al., 2015; Van Hede et al., 2017; Wakita et al., 2010; Wellenstein et al., 2019). How these cells react to tumor-derived factors beyond IL-17A, however, is largely unknown. Therefore, we compared the transcriptome data of Cluster 1 from WT mice with Clusters 1-6 from tumor-bearing KB1P mice to determine which genes are differentially regulated in CD27^—^ γδ T cells by mammary tumors. A total of 96 genes, including *Il17a*, were upregulated in cells from tumor-bearing KB1P mice when compared with cells from WT mice, while 72 genes were downregulated **(Figure 6A; Table 5)**. Among the upregulated genes, *5830411N06Rik* (SCART2), *Tcrg-C2* and *Trgv2* were featured, indicating that Vγ4^+^ cells were more abundant in the lungs of tumor-bearing mice than tumor-free mice, which was also noted by flow cytometry (**Figure 4B**). Other upregulated genes included *Il17f, Havcr2* (TIM-3), *Areg* (amphiregulin), *Irf4*, *Ly6a* (SCA-1), and *Tnfsf8* (CD30L) (**Figure 6A, B**). IL-17F, AREG, IRF4 and SCA-1 are frequently expressed by IL-17A- producing γδ T cells (Jin et al., 2019; Tan et al., 2019; Wang et al., 2021), and CD30L plays a role in maintenance of these cells (Sun et al., 2013). TIM-3 is a co-inhibitory molecule often expressed by CD8^+^ T cells or dendritic cells in the tumor microenvironment (de Mingo Pulido et al., 2021; Dixon et al., 2021; Wolf et al., 2020). We observed that *Il17f* and *Havcr2* expression were enriched in Cluster 5, where Vγ4^+^ cells were localized, while *Tnfsf8* was absent from Cluster 5 and the other genes were evenly distributed across Clusters 1-6 (**Figure 6B**).

**Figure 6:**
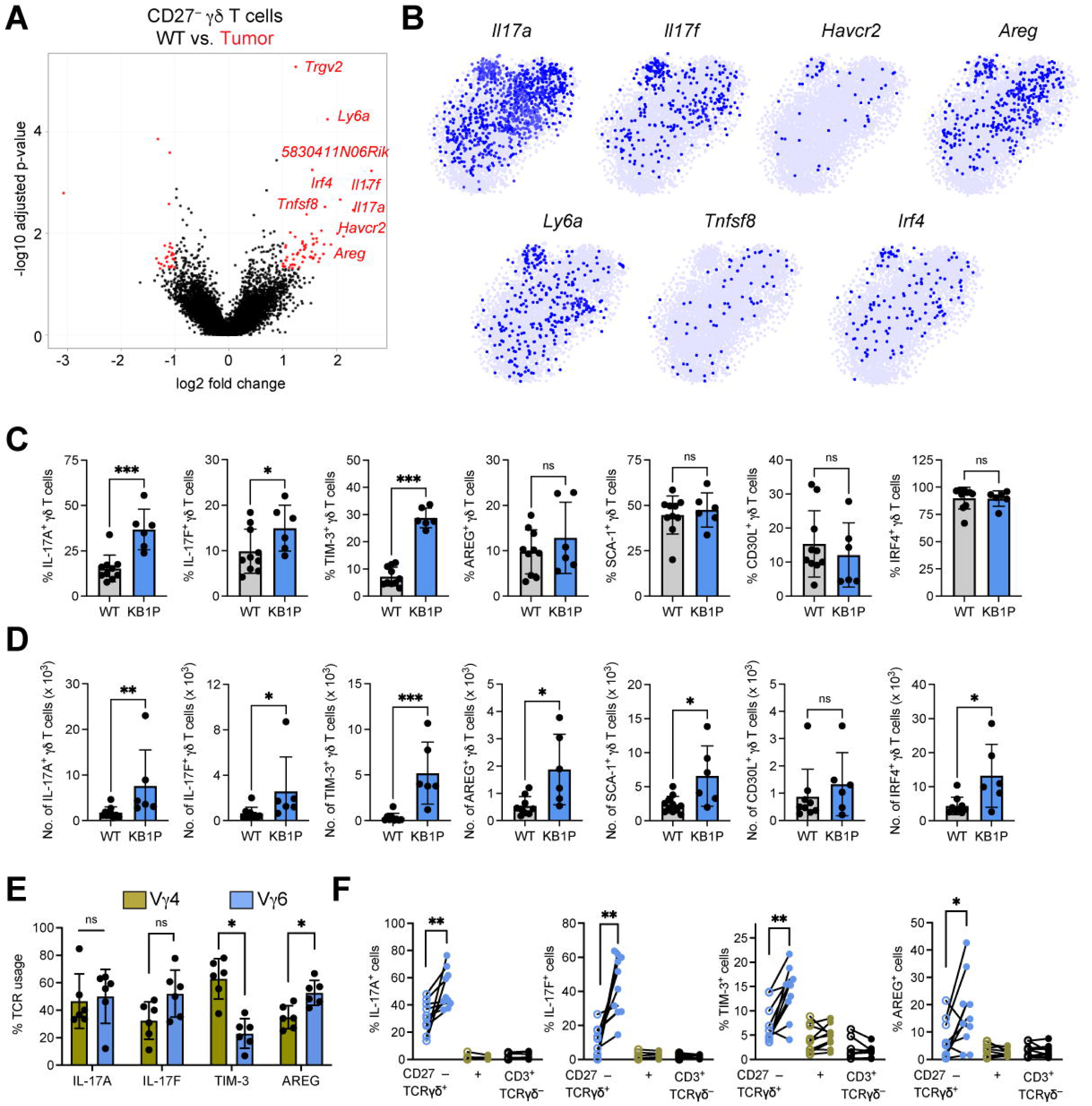
Tumor-derived IL-1β and IL-23 drive gene expression and proliferation in lung γδ T cells. **(A)** The transcriptome of CD27^—^ γδ T cells from KB1P Clusters 1-6 (Figure 4C) was compared to FVB/n Cluster 1 (Figure 1A) and represented as a volcano plot. Genes up- or down-regulated in γδ T cells from KB1P Clusters 1-6 are denoted in red. **(B)** Feature plots in tSNE map of indicated genes. **(C, D)** Single cell suspensions from lung of WT and tumor- bearing KB1P mice were stimulated for 3 hours with PMA, ionomycin and Brefeldin A. Cells were stained with antibodies against CD3, TCRδ and CD27 to identify CD27^—^ γδ T cells by flow cytometry. The proportion of cells expressing the indicated molecules (C) and the absolute number of cells (D) is represented graphically. **(E)** The frequency of IL-17A^+^, IL- 17F^+^, AREG^+^ and TIM-3^+^ CD27^—^ γδ T cells expressing Vγ1, Vγ4, or Vγ6 TCR chains. **(F)** CD3^+^ T cells were isolated from the lungs of FVB/n mice and stimulated with recombinant IL- 1β and IL-23 for 15 hours. Cells were stimulated for 3 hours with PMA, ionomycin and Brefeldin A. Cells were stained with antibodies against CD3, TCRδ and CD27 to distinguish CD27^—^ γδ T cells, CD27^+^ γδ T cells and TCRδ^—^ cells by flow cytometry. The proportion of cells expressing IL-17A, IL-17F, TIM-3 or AREG is shown. Each dot represents one mouse. Data are presented as mean ± SD. \**p* < 0.05, \*\**p* < 0.01, \*\*\**p* < 0.001 as determined by Mann-Whitney U test or Kruskal-Wallis test followed by Dunn’s posthoc test.

We validated the upregulation of these genes at protein level in γδ T cells from the lungs of WT and KB1P tumor-bearing mice by flow cytometry. The frequency of IL-17A-, IL- 17F- and TIM-3-expressing γδ T cells was greater in the lungs of mammary tumor-bearing KB1P mice than WT mice, but the frequency of AREG-, SCA-1-, CD30L- and IRF-4- expressing γδ T cells remained unchanged between groups (**Figure 6C; Supplemental Figure 5**). We observed an increase in the absolute number of IL-17A^+^, IL-17F^+^, TIM-3^+^, AREG^+^, SCA-1^+^ and IRF^+^, but not CD30L^+^, lung γδ T cells in tumor-bearing KB1P mice (**Figure 6D**), which might be explained by the fact that CD27^—^ γδ T cells expand in KB1P mice (**Figure 4A**). Having observed *Il17f and Havcr2* mRNA expression localized to specific γδ T cell subsets (**Figures 5E, 6B**), we investigated whether there was a difference in protein expression of IL-17A, IL-17F, TIM-3 and AREG between lung γδ T cell subsets. After gating on CD27^—^ γδ T cells from KB1P tumor-bearing mice that were either IL-17A^+^ or IL-17F^+^, we found that both Vγ4^+^ and Vγ6^+^ cells are the main producers of IL-17A and IL-17F. By contrast, Vγ4^+^ cells were the main producers of TIM-3 and Vγ6^+^ cells were the main producers of AREG (**Figure 6E**). These data indicate that IL-17A, IL-17F and TIM-3 are specifically induced by mammary tumors, whereas mRNA up-regulation of the other molecules is disconnected from their regulation at the protein level.

Next, we determined how IL-17A, IL-17F, TIM-3 and AREG are regulated in CD27^—^ γδ T cells. In tumor-associated lungs, the systemic increase in IL-1β and IL-23 induces CD27^—^ γδ T cells to expand and rapidly produce cytokines (Coffelt et al., 2015; Jin et al., 2019). To recapitulate the systemic increase of IL-1β and IL-23 in mammary tumor-bearing mice, we stimulated CD3^+^ T cells isolated from lungs of WT mice *in vitro* with these cytokines and examined tumor-associated protein expression in CD27^—^ γδ T cells. This analysis revealed that IL-1β/IL-23 treatment increases expression of IL-17A, IL-17F, TIM-3 and AREG **(Figure 6F)**. These increases were specific to the CD27^—^ γδ T cell population, as CD27^+^ γδ T cells and CD3^+^γδTCR^—^ T cells did not respond to IL-1β/IL-23 stimulation **(Figure 6F)**. These findings indicate that the pro-inflammatory cytokines IL-1β and IL-23 drive the phenotype of pro-tumorigenic Vγ4^+^ and Vγ6^+^ T cells in the lung.

### T cell checkpoint inhibitor immunotherapy increases IL-17A expression by γδ T cells in mammary tumor-bearing mice

The data reported above show that ICOS and PD-1 are constitutively expressed by lung Vγ6^+^ cells and TIM-3 is inducibly expressed by lung Vγ4^+^ cells in response to mammary tumors. Having established the differential expression of these co-inhibitory and co-stimulatory molecules on lung γδ T cells, the impact of PD-1 and TIM-3 inhibition as well as ICOS stimulation on lung γδ T cells was assessed in the orthotopic transplant KB1P model. We performed a short-term treatment experiment over the course of 4 days to mitigate the confounding effects of these antibodies on other T cell subsets. KB1P tumor fragments were transplanted into syngeneic mice and allowed to grow to 1 cm. After this, mammary tumor-bearing mice received daily doses of anti-ICOS, anti-PD-1 or anti-TIM3 over 3 days **(Figure 7A)**. Tumor growth increased in control, anti-PD-1- and anti-TIM-3-treated mice, whereas tumor growth was prevented in anti-ICOS-treated mice (**Figure 7B**). Analysis of lung γδ T cells revealed that inhibition of PD-1 and TIM-3 increases IL-17A expression by two-fold in these cells when compared to control treatment **(Figure 7C**). The ICOS agonistic antibody failed to influence IL-17A expression over controls. Moreover, the effect of blocking PD-1 and TIM-3 signaling on γδ T cells was limited to IL-17A, as IL-17F, AREG and TIM-3 expression were not changed by these treatment modalities **(Figure 7C)**. The absolute numbers of IL-17A-, IL-17F, TIM-3 and AREG-producing γδ T cells remained the same between groups **(Figure 7D)**. These data indicate that inhibition of PD-1 and TIM-3 signaling results in the specific upregulation of IL-17A by γδ T cells in the lung pre-metastatic niche of mammary tumor-bearing mice.

**Figure 7:**
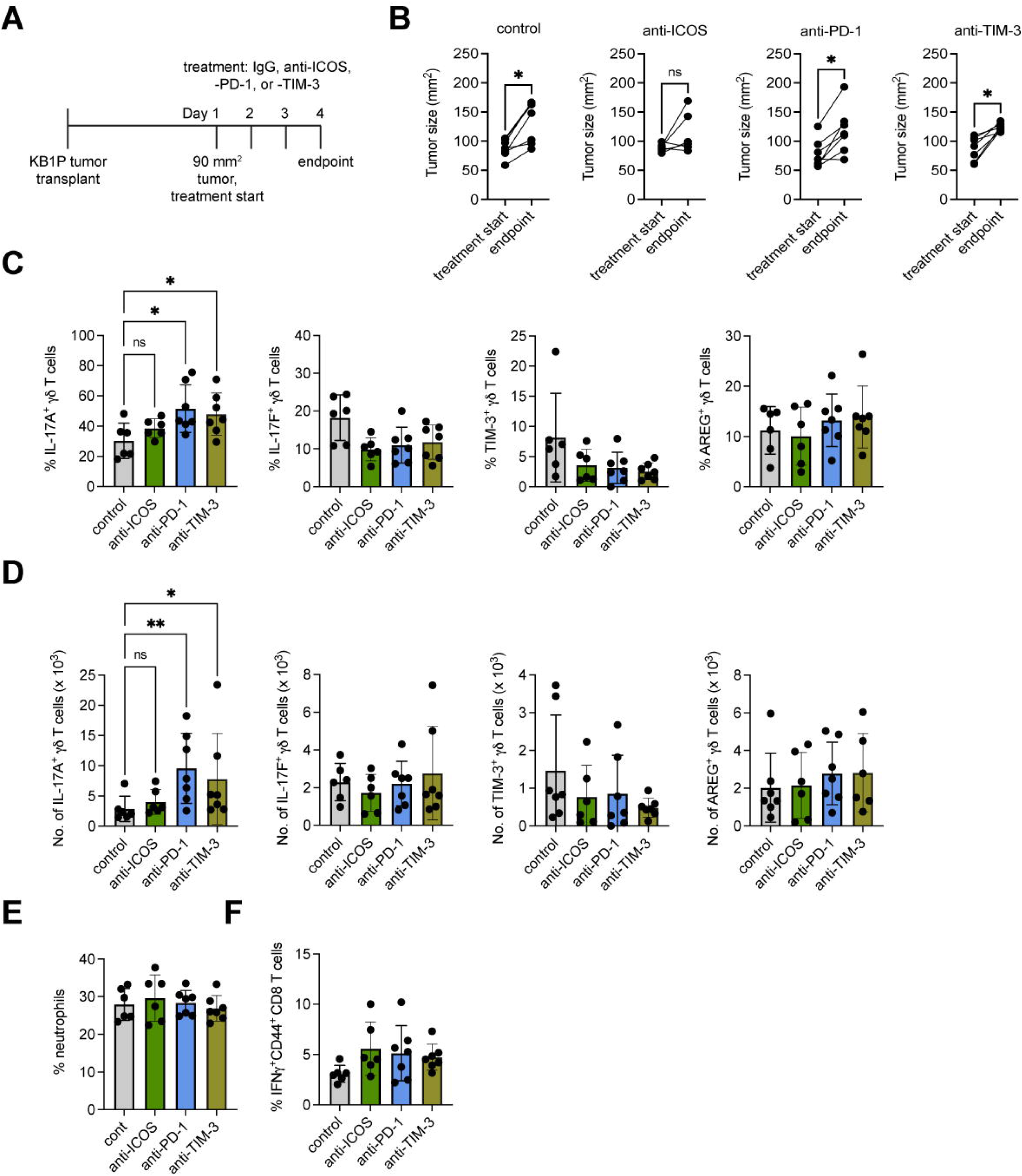
Anti-PD-1 and anti-TIM-3 immunotherapy increases IL-17A expression by lung CD27^—^ γδ T cells in mammary tumor-bearing mice. **(A)** Schematic of experimental procedure for orthotopic transplantation of KB1P tumor fragments into FVB/n mice and treatment with anti-PD1, anti-ICOS, anti-TIM-3, or isotype control. **(B)** Graphic representation of tumor growth at start of treatment and experimental endpoint in mice treated as indicated. **(C, D)** Single cell suspensions from lung of tumor- bearing KB1P mice were stimulated for 3 hours with PMA, ionomycin and Brefeldin A. Cells were stained with antibodies against CD3, TCRδ and CD27 to identify CD27^—^ γδ T cells by flow cytometry. The proportion of cells expressing the indicated molecules (B) and the absolute number of cells (C) is represented graphically. **(E)** Percentage of circulating neutrophils in isotype control, anti-PD-1-, anti-ICOS- and anti-TIM-3-treated tumor-bearing mice as measured by IDEXX ProCyte hematology analyzer. **(F)** Percentage of CD44^+^IFNγ^+^CD8^+^ T cells in lungs of isotype control, anti-PD-1-, anti-ICOS- and anti-TIM-3- treated tumor-bearing mice as measured by flow cytometry. Each dot represents one mouse. Data are presented as mean ± SD. \**p* < 0.05, \*\**p* < 0.01 as determined by Mann-Whitney U test or Kruskal-Wallis test followed by Dunn’s posthoc test.

IL-17A-producing γδ T cells promote metastasis through expansion of neutrophils which in turn inhibit CD8^+^ T cells so that disseminated cancer cells subvert anti-tumor immunity (Coffelt et al., 2015; Wellenstein et al., 2019). Therefore, neutrophil frequency and the phenotype of conventional T cells was investigated in tumor-bearing KB1P mice treated with anti-PD-1, anti-TIM-3 or anti-ICOS, as a systemic readout of increased IL-17A by γδ T cells. However, neutrophil frequency in blood remained the same between treatment groups **(Figure 7E)**, suggesting that the increased IL-17A in this short-term experiment fails to reach a threshold large enough to stimulate granulopoiesis and elevate circulating neutrophils. Expression of CD44 and IFNγ by lung CD8^+^ T cells was also the same between groups **(Figure 7F)**, suggesting that anti-PD-1 and anti-TIM-3 cannot reverse neutrophil-mediated immunosuppression of these CD8^+^ T cells. Overall, acute inhibition of PD-1 and TIM-3 modulated IL-17A in lung γδ T cells without impacting neutrophils or CD8^+^ T cells.

## DISCUSSION

IL-17A-producing γδ T cells within the lung consist mostly of Vγ4^+^ and Vγ6^+^ cells that can function as either protective or pathogenic cells during infection, inflammation and allergy (Faustino et al., 2020; Guo et al., 2018; Misiak et al., 2017; Wang et al., 2021). These cells are also key drivers of lung cancer and pulmonary metastasis (Coffelt et al., 2015; Jin et al., 2019; Kulig et al., 2016). Here, we used scRNAseq with protein validation to provide novel insight into IL-17A-producing γδ T cell subsets in normal and tumor-conditioned lung. Our data show that Vγ6^+^ cells express a phenotype with high degree of similarity to T_rm_ CD8^+^ T cells through production of CXCR6, ICOS, JAML and PD-1, whose expression is stable between tumor-free and tumor-bearing mice. Both Vγ6^+^ and Vγ4^+^ cells rapidly expand and up-regulate various cytokines and growth factors in response to tumor-derived factors, such as IL-1β and IL-23. These cytokines induce TIM-3 specifically on Vγ4^+^ cells in tumor-bearing mice. We found that IL-17A in Vγ6^+^ cells is regulated by PD-1 signaling, while IL-17A in Vγ4^+^ cells is regulated by TIM-3, highlighting the differential control of these discrete subsets by distinct co-inhibitory molecules.

The expression and function of co-stimulatory and co-inhibitory molecules on IL-17A- producing γδ T cells has received little attention. In agreement with our data, other studies have reported high levels of ICOS and PD-1 expression on Vγ6^+^ cells from the uterus and skin (Monin et al., 2020; Tan et al., 2019), suggesting that these two molecules are common features of Vγ6^+^ cells in any tissue. PD-1-deficient mice exhibit elevated levels of IL-17A from γδ T cells and greater disease severity of imiquimod-driven psoriasis (Imai et al., 2015; Kim et al., 2016), which accords well with our observations. We show that PD-1 signaling can suppress IL-17A expression in Vγ6^+^ cells; however, the role of ICOS on these cells is unclear. The ICOS/ICOSL axis is critical for development of IL-17A-producing γδ T cells. ICOS-deficient mice develop increased numbers of Vγ4^+^ cells than wild-type controls, and experimental autoimmune encephalomyelitis (EAE) is more severe in these mice as a result of increased IL-17A levels (Buus et al., 2016; Galicia et al., 2009). Whether the development of Vγ6^+^ cells is affected by loss of ICOS remains unknown. For Vγ4^+^ cells, our data show that TIM-3 expression is positively regulated by the cytokines, IL-1β and IL-23. It seems that TIM- 3 is not the only co-inhibitory receptor that is inducible on Vγ4^+^ cells, as BTLA, CTLA-4 and PD-1 are also increased by IL-1β, IL-23 or IL-7 stimulation (Bekiaris et al., 2013; Kadekar et al., 2020). Taken together, these observations have important implications for the regulation of discrete γδ T cell subsets in the lung during homeostasis and disease.

A question remains about whether PD-1 and TIM-3 regulation of IL-17A expression is mediated through a shared mechanism. PD-1 signaling through SHP-1 and SHP-2 phosphatases disrupts TCR signaling and CD28 co-stimulation via inhibition of LCK and ZAP70 activation of PI3K/AKT and MAPK pathways (Sharpe and Pauken, 2018). However, in IL-17A-producing γδ T cells, release of cytokines is generally considered independent of TCR activation. PD-1 signaling is known to activate the transcription factor, BATF, to impair CD8^+^ T cell proliferation (Quigley et al., 2010), and BATF-deficient IL-17A-producing γδ T cells are hyper-proliferative (Barros-Martins et al., 2016; McKenzie et al., 2017). Therefore, it is tempting to speculate that PD-1 activation on Vγ6^+^ cells induces BATF-mediated suppression of cell proliferation and IL-17A expression. TIM-3 binds an adaptor protein called BAT3 (or BAG6) that negatively regulates its function (Rangachari et al., 2012; Wolf et al., 2020). Activation of TIM-3 by Galetin-9 or another ligand releases its interaction with BAT3, which leads to suppression of mTORC2 and AKT phosphorylation, as well as nuclear localization of the transcriptional factor, FOXO1 (Zhu et al., 2021). FOXO1 is an inhibitor of the IL-17A master regulator, RORγt (Laine et al., 2015). Thus, blockade of TIM-3 may encourage its association with BAT3, allowing phosphorylation of mTORC2, AKT and FOXO1 that relocalizes FOXO1 to the cytoplasm. In this way, RORγt-mediated transcription of *Il17a* mRNA would go forward without interference from FOXO1. Alternatively, TIM-3 binding to the SRC tyrosine kinase family members, LCK and FYN, may impact on RORγt signaling to mediate suppression of IL-17A secretion (Lee et al., 2011; Rangachari et al., 2012; Ueda et al., 2012; Wolf et al., 2020). Further experimentation is required to fully elucidate the mechanism of PD-1 and TIM-3 suppression of IL-17A in γδ T cells.

In our scRNAseq data, we observed that *Tmem176a* and *Tmem176b* mRNA are highly expressed by lung CD27^—^ γδ T cells. Previous reports on skin γδ T cells have demonstrated that these ion channels are exclusively expressed by Vγ6^+^ cells (Tan et al., 2019), and *Tmem176a* and *Tmem176b* are strongly expressed by other RORγt-producing cells (Drujont et al., 2016). Deficiency of *Tmem176b* in mice reduces imiquimod-induced, IL- 17A-mediated psoriasis (Drujont et al., 2016), suggesting its importance in the regulation of IL-17A-producing cells. In addition, targeting TMEM176B is important for the success of checkpoint inhibitor immunotherapy, as evidenced by the response rates of tumor-bearing mice that lack *Tmem176b* to anti-PD-1 and anti-CTLA-4 antibodies (Segovia et al., 2019). Furthermore, a TMEM176B small molecule inhibitor can restore anti-CTLA-4 and anti-PD-1 anti-tumor responses by promoting cytotoxic CD8^+^ T cell effector function (Segovia et al., 2019). It would be interesting to establish whether targeting TMEM176A and TMEM176B on IL-17A-producing γδ T cells could be used in combination with anti-PD-1 or anti-TIM-3 immunotherapy to reduce IL-17A expression by Vγ6^+^ and Vγ4^+^ cells and control cancer progression.

There is evidence from clinical studies to suggest that IL-17A signaling undermines immune checkpoint inhibitors, contributes to immunotherapy resistance and promotes the development of adverse autoimmune events in cancer patients (Kang et al., 2021). As such, modulation of PD-1 and TIM-3 in cancer patients and the impact of inhibiting these molecules on γδ T cells is a point for consideration, as these immunotherapy drugs may deleteriously increase IL-17A expression and immunosuppressive neutrophil activation. Indeed, immune checkpoint inhibitor non-responding melanomas contain a greater proportion of a γδ T cell subset than responding melanomas, and a gene signature derived from these γδ T cells can predict response to immune checkpoint inhibitors (Xiong et al., 2020). The effect of immune checkpoint inhibitors on IL-17A is not limited to γδ T cells, as human CD4^+^ T cells increase IL-17A after anti-PD-1 exposure (Bandaru et al., 2014) and IL-17A-producing cells in general are associated with anti-PD-1 resistance in melanoma, lung and colorectal cancer (Gopalakrishnan et al., 2018; Li et al., 2021; Llosa et al., 2019; Peng et al., 2021). In agreement with our data, anti-PD-1 treatment of an autochthonous lung cancer mouse model increases IL-17A-producing γδ T cells and CD4^+^ T cells, which correlates with suppression of cytotoxic CD8^+^ T cells (Li et al., 2021). Moreover, IL-17A-overexpressing lung tumors are resistant to anti-PD-1 therapy (Akbay et al., 2017). However, inhibition of IL-17A in mice bearing KRAS-mutant lung tumors or transplantable colon cancer cell lines has provided proof-of-principle that targeting IL-17A in combination with anti-PD-1 is a viable strategy for controlling tumor growth (Li et al., 2021; Liu et al., 2021; Peng et al., 2021). Anti-PD-1 immunotherapy, although effective against a minority of melanomas, can also exacerbate psoriasis and colitis (Tanaka et al., 2017), but IL-17A blockade may reverse these toxicities (Esfahani and Miller, 2017; Johnson et al., 2019). Taken together, our observations reported herein and studies in the literature suggest that targeting IL-17A in combination with immune checkpoint inhibitors may thwart resistance mechanisms and lessen adverse autoimmune events in cancer patients.

Collectively, our data offers novel insights into the unique phenotype of specialized lung γδ T cell subsets. Our data demonstrate that lung γδ T cell subsets are distinctly regulated by different T cell checkpoint molecules in both homeostasis and cancer. These data have implications for the role of γδ T cells in other lung pathologies and infection.

## ACKNOWLEDGEMENTS

We thank Catherine Winchester and all Coffelt lab members for critical discussion, contribution to the development of this work and helpful comments on the manuscript. We thank Jos Jonkers for the mammary cancer mouse model. We thank Shinya Hatano and Yasunobu Yoshikai for Vγ6 hybridoma. We would like to thank the Core Services and Advanced Technologies at the Cancer Research UK Beatson Institute, with particular thanks to the Biological Services Unit.

## AUTHOR CONTRIBUTIONS

SCE and SBC: conceptualization. SCE, AH, TG, WHMH, RW, AK, RS, KB, NB, KK and SBC: data acquisition, analysis and visualization. SCE and SBC: wrote the manuscript. All authors: review and editing manuscript. KB, CM and SBC: supervision. SBC: funding acquisition.

## FIGURE LEGENDS

**Supplemental Figure 1: Gating strategy for sorting γδ T cells.**

Single cell suspensions of lung cells were stained with antibodies against CD3 and TCRδ. DAPI was used to exclude dead cells. Flow cytometry plots are depicted showing doublet exclusion, live cells, T cells and γδ T cells. Numbers represent the frequency of cells in each gate.

**Supplemental Figure 2: Differentially expressed genes in Cluster 1 and Cluster 2 of γδ T cells from WT FVB/n mice.**

**(A, B, C)** Feature plots in tSNE map of indicated genes from FVB/n mice.

**Supplemental Figure 3: Lung CD27^—^ γδ T cells in C57BL/6 mice express a tissue-resident memory phenotype.**

Single cell suspensions of lung from C57BL/6J mice were stained with antibodies against CD3, TCRδ, CD27 and the indicated molecules. Cells were analyzed by flow cytometry. Graphic representation of percentage positive cells for each indicated molecule in CD27^+^ and CD27^—^ γδ T cells from lung. Each dot represents one mouse. Data are presented as mean ± SD. \**p* < 0.05 as determined by Mann-Whitney U test.

**Supplemental Figure 4: Pathway analysis of scRNAseq clusters identified from γδ T cells in mammary tumor-bearing mice.**

Differentially expressed genes for each individual cluster were analyzed by REACTOME pathway analysis. The top 5 pathways characterizing each cluster is shown.

**Supplemental Figure 5: Expression of IL-17A, IL-17F, TIM-3, AREG, SCA-1 and CD30L by lung CD27^—^ γδ T cells.**

Flow cytometry plots of staining for indicated molecules in lung CD27^—^ γδ T cells from WT and tumor-bearing KB1P mice. Single cell suspensions of lung were stimulated for 3 hours with PMA, ionomycin and Brefeldin A, and stained for extracellular and intracellular markers and analysed by flow cytometry.

