## Supplementary figures and images for "Single-cell analysis uncovers differential regulation of lung γδ T cell subsets by the co-inhibitory molecules, PD-1 and TIM-3"

### Supplemental Figure 1

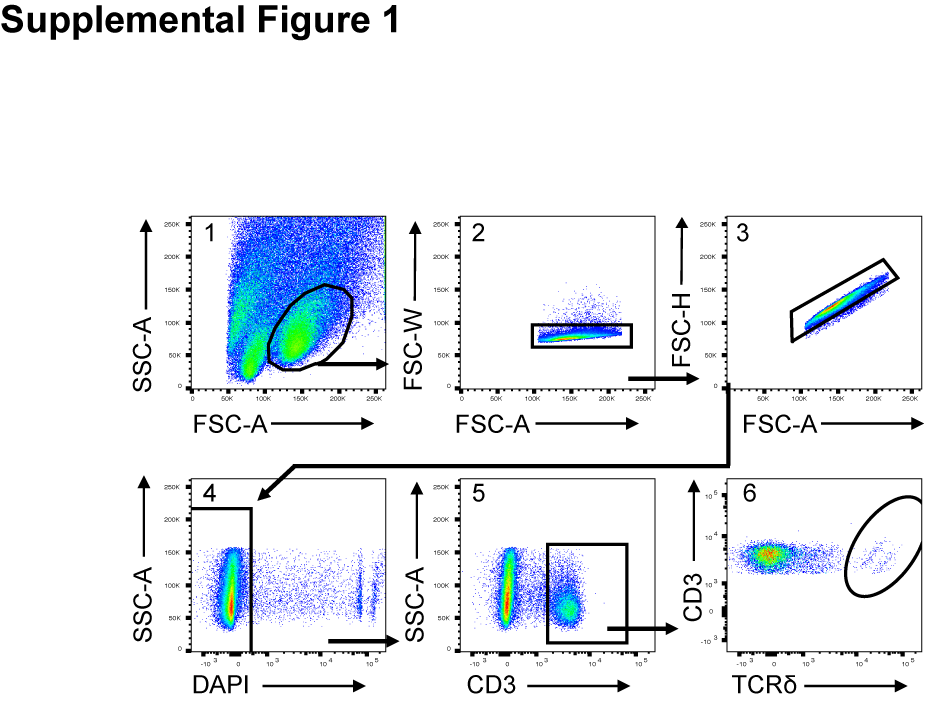

### Supplemental Figure 2

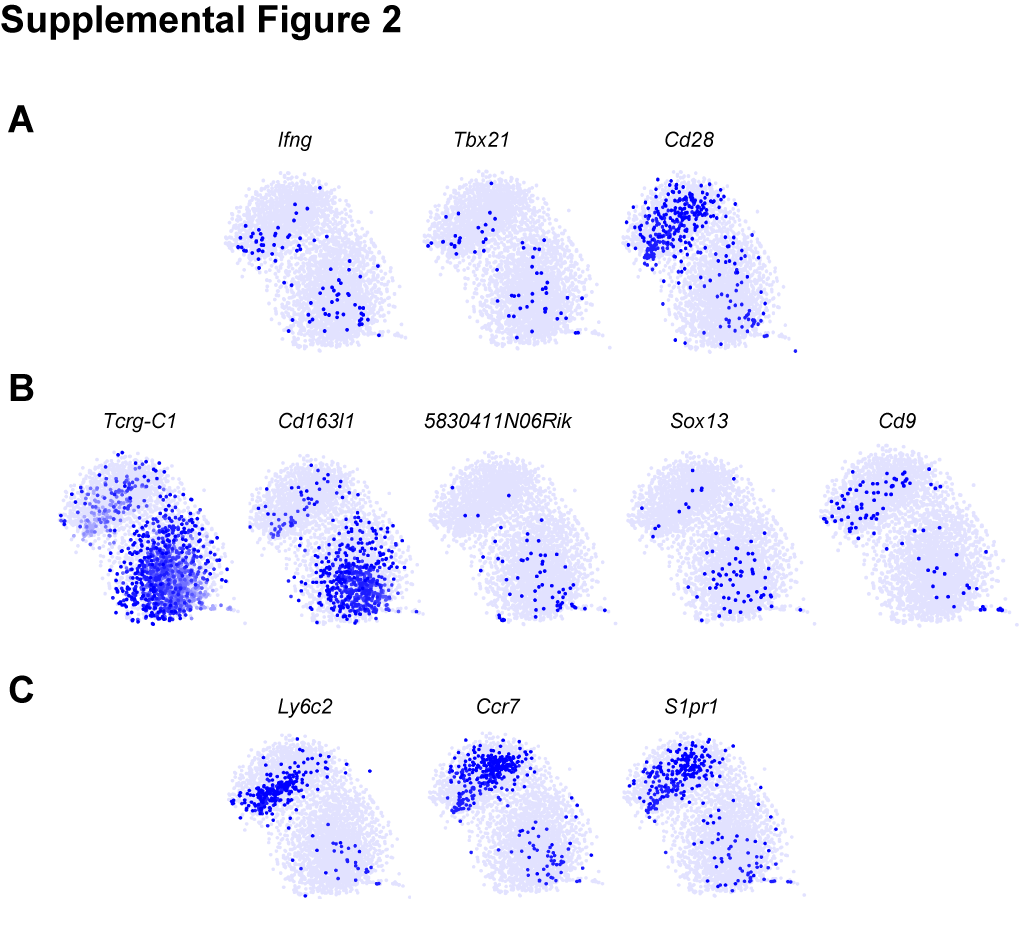

### Supplemental Figure 3

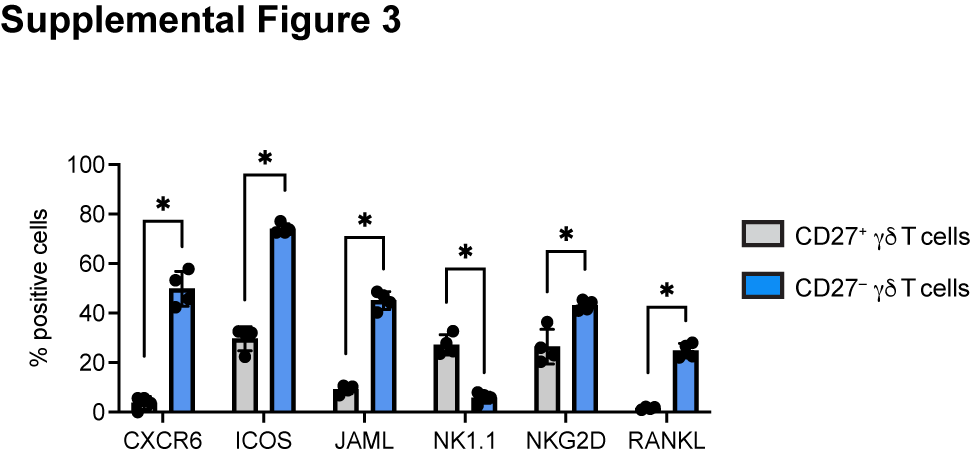

### Supplemental Figure 4

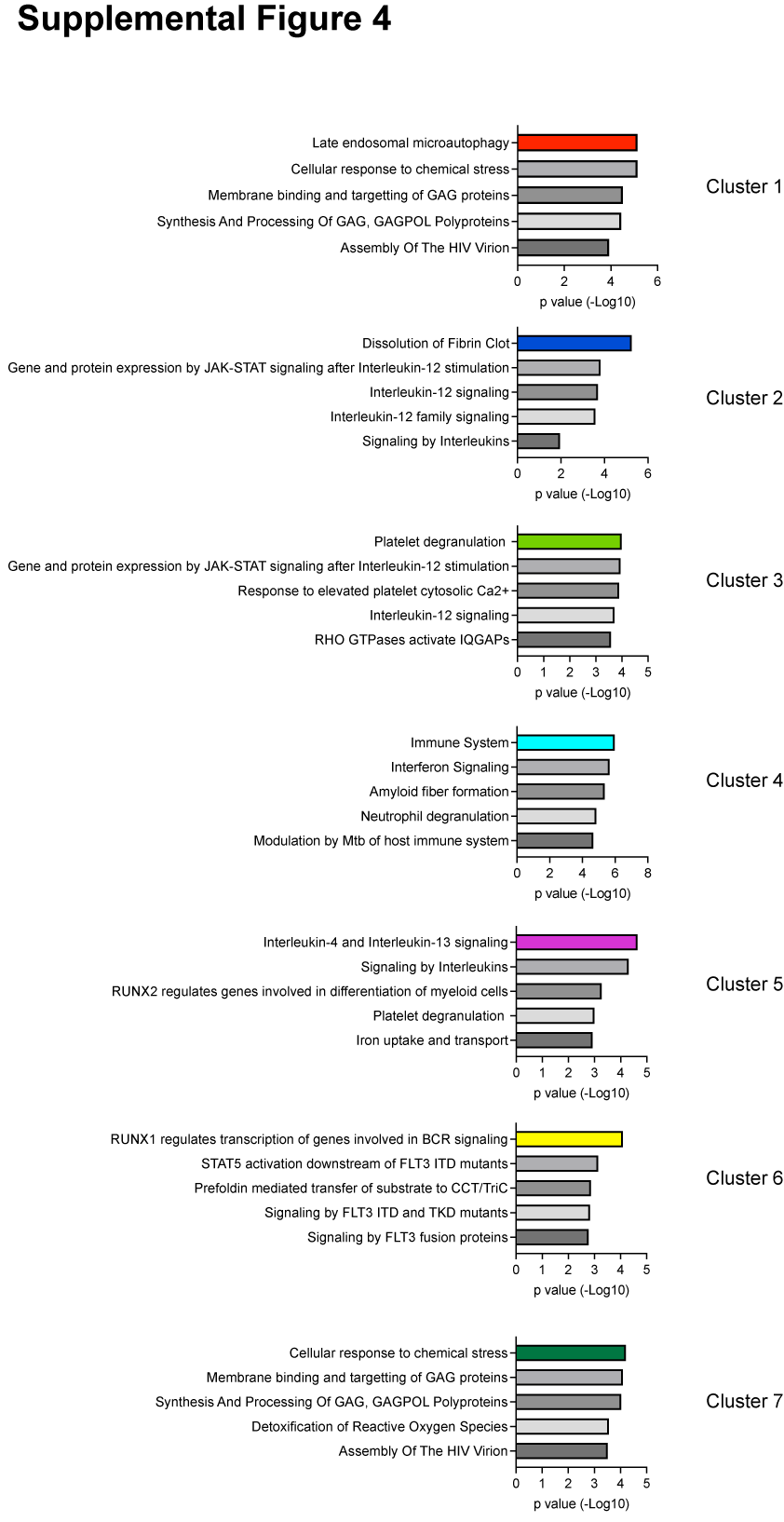

### Supplemental Figure 5

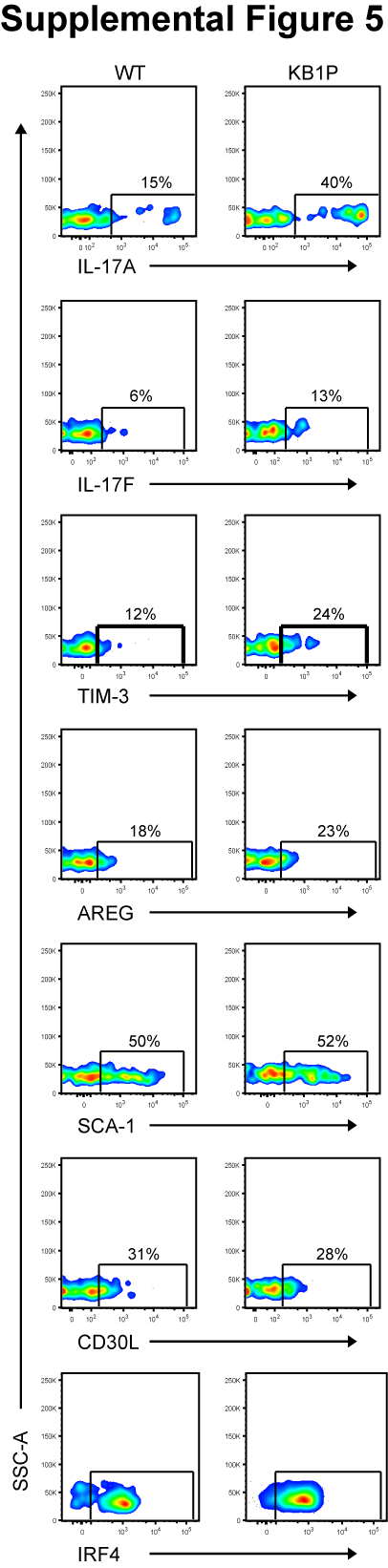
